# Profound CD4+ T-Cell Reprogramming by Melphalan-Driven Oxidative Stress in High-Risk Multiple Myeloma

**DOI:** 10.64898/2026.06.08.730835

**Authors:** Mara John, Nazia Afrin, Xiang Zhou, Emilia Stanojkovska, Hanna Fischer, Maximilian J. Krämer, Werner Schmitz, Lars Grundheber, Moutaz Helal, Arpa Aintablian, Silvia Nerreter, Cornelia Vogt, Shilpa Kurian, Angelika Wolf, Sarah Wiethe, Maximilian J. Steinhardt, Benedict M. Engel, Anni Hofmann, Marc Schmalzing, Michael Hudecek, Hermann Einsele, K. Martin Kortüm, Leo Rasche, Angela Riedel

**Author notes:** Corresponding authors: Angela Riedel; Leo Rasche. These authors contributed equally (Mara John, Nazia Afrin, Xiang Zhou). These authors are shared last authors (Leo Rasche, Angela Riedel).

## Abstract

T cell-based immunotherapies have become central to the treatment of multiple myeloma (MM), yet their efficacy depends on the functionality of endogenous T cells. How cumulative treatment exposure, particularly high-dose melphalan, together with disease-intrinsic high-risk features shapes T-cell composition and immune competence remains incompletely understood. Here, we analyzed T cell composition and function in bone marrow (BM) and peripheral blood (PB) samples from MM patients across different stages of their treatment journey using flow cytometry (BM, n=162; PB, n=1,733), single-cell RNA sequencing (n=19), and cytotoxicity assays (n=20). We reveal reduced overall T cell frequencies and CD4+/CD8+ T cell ratio, associated with lines of therapy and driven in part by depletion of naïve CD4+ T cells in gene-expression defined high risk (HR) disease. Among therapeutic agents, melphalan exerted the strongest effects on T cell populations and induced pronounced redox stress in both T cells and myeloma cell lines. This oxidative stress signature was enriched in HR patients and was reversible with N-acetyl-L-cysteine treatment. Together, these findings identify immune dysregulation as a defining feature of HR MM that extends beyond tumor-intrinsic genomic alterations and is further shaped by treatment-induced remodeling of the BM microenvironment. Given the association between higher CD4+ T cell numbers and improved CAR-T cell outcomes, our data highlight the translational importance of treatment sequencing, particularly in HR MM.

**One Sentence Summary:** Melphalan-induced redox stress depletes CD4+ naïve T cells, particularly in patients with high-risk multiple myeloma.

## INTRODUCTION

Multiple myeloma (MM) has seen a revolution in immunotherapies with many patients benefitting from T cell-based therapies, such as chimeric antigen receptor (CAR) T cells and bispecific antibodies (bsAbs). Most of these therapies have been implemented to treat refractory or relapsed MM (RRMM). However, they rely heavily on functional endogenous T cells. Therefore, it is surprising that the effect of different prior therapies on patients’ T cells and their functionality has not yet been thoroughly studied. The efficacy of T cell-based therapies was classically believed to rely on CD8+ T cell mediated tumor cell killing. However, CD4+ T cells have emerged as key players in the outcomes of immunotherapy and anti-tumor immunity *(1–3)*. In an elegant study, Kruse et al. *(4)* demonstrated that CD4+ T cells can eradicate MHC-deficient tumor cells, whereas CD8+ T cells cannot. Moreover, a higher CD4+ T cell proportion in leukapheresis used for CAR-T generation in MM is correlating with better clinical response *(5)*. In particular, it has been shown that cytotoxic CD4+ T cells in CAR-T cell products drive tumor killing and durable disease control *(6)*.

Combinatorial high-dose chemotherapy, which combines treatment schedules with agents such as vincristine, epirubicin, dexamethasone, thalidomide, bendamustine, prednisone, cisplatin, etoposide and melphalan, is known to alter T cell subsets, with effects lasting for over two years after treatment. However, more detailed analyses were missing *(7)*. Melphalan, a DNA alkylating agent, reduced the expansion and transduction of T cells used for CAR-T cell therapy in MM *(8)*. Furthermore, in B-cell non-Hodgkin lymphoma, it was observed that prior lines of treatment negatively impacted the CAR-T cell product, resulting in reduced killing capacity of T cells from treated patients compared to T cells from untreated patients *(9)*.

High-risk (HR) MM, a disease category characterized by early relapse and inferior survival outcomes, may further impose profound alterations in T cell composition and immune competence. In particular, HR MM patients are at a higher risk of non-relapse mortality, which is mostly due to opportunistic infections *(10)*. In addition, newly diagnosed HR patients have a significantly increased risk to suffer from severe infections within the first 4 months after diagnosis *(11)*. HR patients who experienced a relapse within 24 months and then received CAR-T cells in later lines of therapy (five or more) had worse overall survival compared to HR patients who experienced a longer period until relapse *(12)*.

In this study, we thus investigated the effect of commonly used MM therapies on the T cell compartment in MM patients in the peripheral blood (PB) and bone marrow (BM) in dependency of gene-array defined risk status. We demonstrate that CD4+ T cell counts decrease specifically after melphalan treatment, and that this decrease is associated with poorer progression-free and overall survival. Additionally, we found a more pronounced reduction in CD4+ T cells in the PB and BM of HR patients induced by treatment. Treatment also reduced the *in vitro* killing capacity of T cells isolated from the BM of HR patients. Furthermore, treatment impacted T cell subpopulations; specifically, CD4+ naïve T cells were depleted in treated HR MM patients, which was associated with an upregulation of apoptosis markers. Finally, melphalan-induced redox stress reduced naïve T cells *in vitro* and was attenuated by N-acetyl-L-cysteine supplementation. Taken together, our data call into question the use of melphalan treatment prior to T cell-based immunotherapies, especially for HR MM patients.

## RESULTS

### CD4 T cell counts decline in melphalan treated patients and predict patient survival

To investigate the impact of treatment and risk status on T cell populations, we collected PB (n=1733) and BM (n=162) samples from MM patients undergoing standard treatment in our clinic (Fig. 1A). All samples were profiled using flow cytometry and markers for immune cells (CD45), as well as T cells (CD3) and their conventional CD8 and CD4 subsets. PB T cell populations of RRMM patients were then analyzed in dependency of the treatment they received covering MM relevant therapies, such as high-dose melphalan, bortezomib, carfilzomib, lenalidomide, pomalidomide, elotuzumab, daratumumab, and BCMA-targeting antibody drug conjugate belantamab mafodotin. This revealed that overall many therapies affected T cell numbers and percentages (Fig. 1B, fig. S1, A to G, table 1), mostly inducing a reduction in CD4+ T cells, both in total numbers and percentages (% of CD3), and an elevation of CD8+ T cells, both in total numbers and percentages (% of CD3), in comparison to patients not receiving that specific therapy. Still, only carfilzomib, a proteasome inhibitor, and high-dose melphalan, a DNA alkylating agent, were found to be statistically significant in reducing CD4+ T cells of overall CD3+ T cells (%) and total numbers (median reduction 14.29% in carfilzomib and 44.83% in melphalan in the case of proportion (%) and 34.77% in carfilzomib and 43.34% in melphalan in total numbers; p < 0.0001 in all cases), while bortezomib, another proteasome inhibitor, showed no significant difference (Fig. 1, C to E). In addition, patients treated with melphalan exhibited a concomitant decrease in the frequency of total T cells (CD3+) among CD45+ immune cells, while total T cells were unchanged, in comparison to patients not receiving high-dose melphalan (fig. S1, H and I). The same patient group had increased proportions and absolute numbers of CD8+ T cells (fig. S1, J and K). In summary, high-dose melphalan demonstrated the severest depletion of the proportion and absolute number of CD4+ T cells in PB (median reduction 44.83% in the case of proportion and 43.34% in absolute number; p < 0.0001 in both cases) (Fig. 1E).

**Fig. 1.**
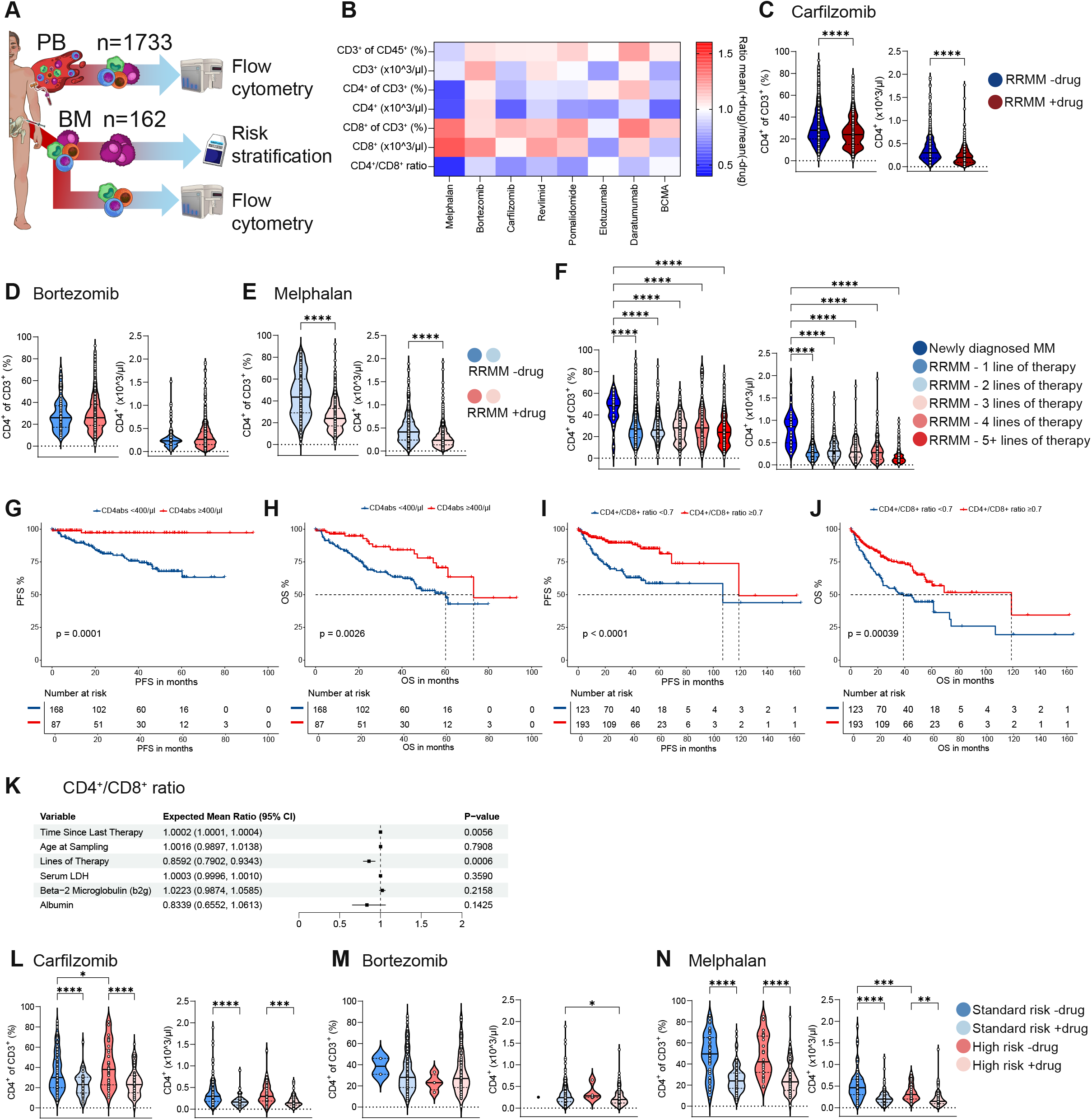
Therapy induces loss in CD4+ T cell population and is linked with worse patients’ survival. (**A**) Experimental setup for flow cytometry analysis from peripheral blood (PB) and bone marrow (BM) and risk stratification from plasma cells (PC) from BM. (**B**) Heatmap of effect of drugs on T cell populations in the PB of relapsed/refractory multiple myeloma (RRMM) patients analyzed by flow cytometry. (**C** to **E**) Quantification of flow cytometry results from PB of RRMM patients. Percentage CD4+ T cells from CD3+ T cells (left panel) and absolute number of CD4+ T cells (right panel) are plotted in regards to carfilzomib (% CD4: −drug n=1031; +drug n=629; absolute CD4+: −drug n=926; +drug n=511) (C), bortezomib (% CD4: −drug n=88; +drug n=1572; absolute CD4+: −drug n=73; +drug n=1364) (D), and melphalan (% CD4: −drug n=320; +drug n=1340; absolute CD4+: −drug n=281; +drug n=1156) (E). Darker colors represent patients who did not receive this drug (RRMM −drug), whereas lighter colors represent patients who received this drug (RRMM +drug). Two-sided unpaired t-test was used for statistical testing. (**F**) Quantification of flow cytometry for percentage of CD4+ T cells of CD3+ T cells (left panel, sample numbers from left to right: n=44; n=686; n=280; n=201; n=174; n=319) and absolute number of CD4+ T cells (right panel, sample numbers from left to right: n=39; n=623; n=236; n=178; n=136; n=264) in the PB of newly diagnosed multiple myeloma (NDMM) and RRMM patients. RRMM patients were separated depending on lines of therapy. Ordinary one-way ANOVA with Fisher’s Least Significant Difference (LSD) as a Post-hoc test was used for statistical testing. (**G** and **H**) Progression-free survival (PFS; G) and overall survival (OS; H) analysis of RRMM patients in dependency of absolute CD4+ T cell count in the PB. Blue < 400 CD4+ T cells/µl PB; red ≥ 400 CD4+ T cells/µl PB. (**I** and **J**) PFS (I) and OS (J) analysis of RRMM patients in dependency of CD4+/CD8+ T cell ratio in the PB. Blue < 0.7 CD4+/CD8+; red ≥ 0.7 CD4+/CD8+. (**K**) Multivariable forest plot visualizing the point estimates (EMR) and 95% confidence intervals (CI) showing the effect of different clinical features on CD4+/CD8+ T cell ratio in the PB. (**L** to **N**) Quantification of flow cytometry results from PB in dependency of risk status and if the patient received the drug. Percentage CD4+ T cells of CD3+ T cells (left panel) and absolute number of CD4+ T cells (right panel) in regards to carfilzomib (% CD4+ (absolute CD4+) SR −drug n=165 (n=152); SR +drug n=54 (n=44); HR −drug n=50 (n=48); HR +drug n=83 (n=64)) (K), bortezomib (% CD4+ (absolute CD4+) SR −drug n=2 (n=1); SR +drug n=217 (n=195); HR −drug n=4 (n=4); HR +drug n=129 (n=108)) (L), and melphalan (% CD4+ (absolute CD4+) SR −drug n=64 (n=61); SR +drug n=155 (n=135); HR −drug n=34 (n=33); HR +drug n=99 (n=79)) (M). Ordinary one-way ANOVA with Fisher’s LSD as a Post-hoc test was used for statistical testing. * p < 0.05, ** p < 0.01, *** p < 0.001, **** p < 0.0001.

CD4+ T cells play an important role in immunotherapy, as well as in the efficacy of CAR-T cells and their production *(5, 13, 14)*. As the decline of CD4+ T cells was seen in many different therapies, e.g. proteasome inhibitor, alkylating agent, immunomodulatory drugs, we wanted to examine if this effect is accumulating over several lines of therapy. Therefore, we compared newly diagnosed MM (NDMM) patients with RRMM patients, separated into different lines of therapy. The depletion of CD4+ T cells was an immediate effect of treatment, while there was no accumulation or recovery of this effect over time, probably attributed to the use of high-dose melphalan in the first line (Fig. 1F). To investigate whether CD4+ T cells play a role not only in immunotherapy but also in overall patient survival, given that a high proportion of patients succumb to infectious complications, we analyzed patient overall and progression-free survival (PFS), stratifying patients based on high (≥400/μL PB) or low (≤400/μL PB) levels of CD4+ T cells in PB, as well as a high (≥0.7) or low (<0.7) ratio of CD4+ to CD8+ T cells (optimal cutoff was defined using surv-cutpoint function *(15)*). Strikingly, this revealed that high CD4+ T cell levels (≥400/μL PB) was linked with longer PFS and overall survival (OS; Fig. 1, G and H). Similarly, patients who have a higher CD4+/CD8+ ratio (≥0.7) showed longer PFS and OS (median of 119 months in both PFS and OS; p < 0.0001 and p < 0.0004 respectively) (Fig. 1, I and J). In general, lines of therapy were negatively associated with a higher CD4+/CD8+ T cell ratio (95% CI, 0.7902 to 0.9343, p < 0.001), while a longer time period between the last treatment and sampling date was positively associated (95% CI, 1.0001 to 1.004; p < 0.01) (Fig. 1K).

Patients with HR MM demonstrate an elevated probability of non-relapse mortality, a phenomenon predominantly attributable to infections, which may be due to an impaired T cell response. To study the combined effect of treatment type and risk status on T cell populations in the BM, we processed BM aspirates of MM patients and CD138+ plasma cells (PC) were used to risk stratify patients into HR and standard-risk (SR) based on the SKY92 profiler (Fig. 1A; *(16)*), whereas the CD138-cells were analyzed via flow cytometry. The SKY92 profiler uses a 92-gene signature to stratify patients into SR or HR with HR being associated with a worse OS. HR patients defined by this SKY92 profiler also show an enrichment for the classical cytogenetic aberrations 1q gain, del(17p), t(4;14), t(14;16), t(14;20) and del(13q) *(16)*. BM T cell populations of RRMM patients were then analyzed in dependency of the treatment they received covering carfilzomib, bortezomib and high-dose melphalan and of their risk status. In both risk status’ we observed a reduction in CD4+ T cells, both in percentages (% of CD3) and in total numbers after treatment with carfilzomib and high-dose melphalan, while bortezomib showed no effect on the proportion in comparison to the same patient groups not receiving the therapy (Fig. 1, L to N, and table 2). However, HR patients showed reduced total CD4+ T cells already in the group not receiving high-dose melphalan (-drug group) in comparison to SR patients, indicating inherent differences of T cells between the two risk groups (Fig. 1M). Additionally, while the high-dose melphalan induced loss of CD3+ T cells was observed in both risk groups after melphalan treatment (fig. S1L), the absolute number of CD3+ T cells was also already depleted in HR patients who did not receive high-dose melphalan compared to SR patients (fig. S1M). The proportion and absolute number of CD8+ T cells increased in the two risk groups in patients receiving high-dose melphalan treatment (fig. S1, N and O). In summary, commonly used MM therapies deplete CD4+ T cells in MM in the PB and in the BM, with the most severe effect of melphalan. Remarkably, the sheer number of CD4+ T cells or the CD4+/CD8+ ratio predicted patient survival, underscoring the indispensable role of viable CD4+ T cells in these patients.

### High-risk patients show more pronounced CD4+ T cell loss and reduced killing capacity after treatment

As we observed differences in the T cells of patients of the two risk groups that may be due to innate differences, we next compared the influence of risk status on T cell populations, independent of treatment. Furthermore, we compared PB and BM to investigate whether these two compartments differ. Notably, HR patients had a higher rate of BM plasma cell infiltration than SR patients in our set (fig. S2A and table 3), which potentially influences the BM T cell compartment.

When analyzing the PB, we found no significant difference of CD3 percentages (% of CD45), as well as numbers of CD3+ T cells, proportion of CD4+ T cells, the ratio of CD4+/CD8+, and the absolute number of CD8+ T cells between SR and HR patients (Fig. 2A and D and fig. S2, B to E). However, the absolute number of CD4+ T cells was reduced in HR patients compared to SR patients (Fig. 2B) and the proportion of CD8+ T cells was increased (% of CD3, Fig. 2C). When we compared NDMM against RRMM in the same cohort, differences in the proportion of CD4+ T cells and other T cell subsets were found in SR and HR patients comparing NDMM and RRMM, with significance only reached in SR. However, the data set was limited due to the small number of patients with NDMM HR (Fig. 2E and fig. S2, F to J). The absolute number of CD4+ T cells, both in SR and HR patients was reduced in RRMM patients compared to NDMM patients, with significance also observed between SR and HR RRMM, indicating a more pronounced phenotype in HR RRMM (Fig. 2F).

**Fig. 2.**
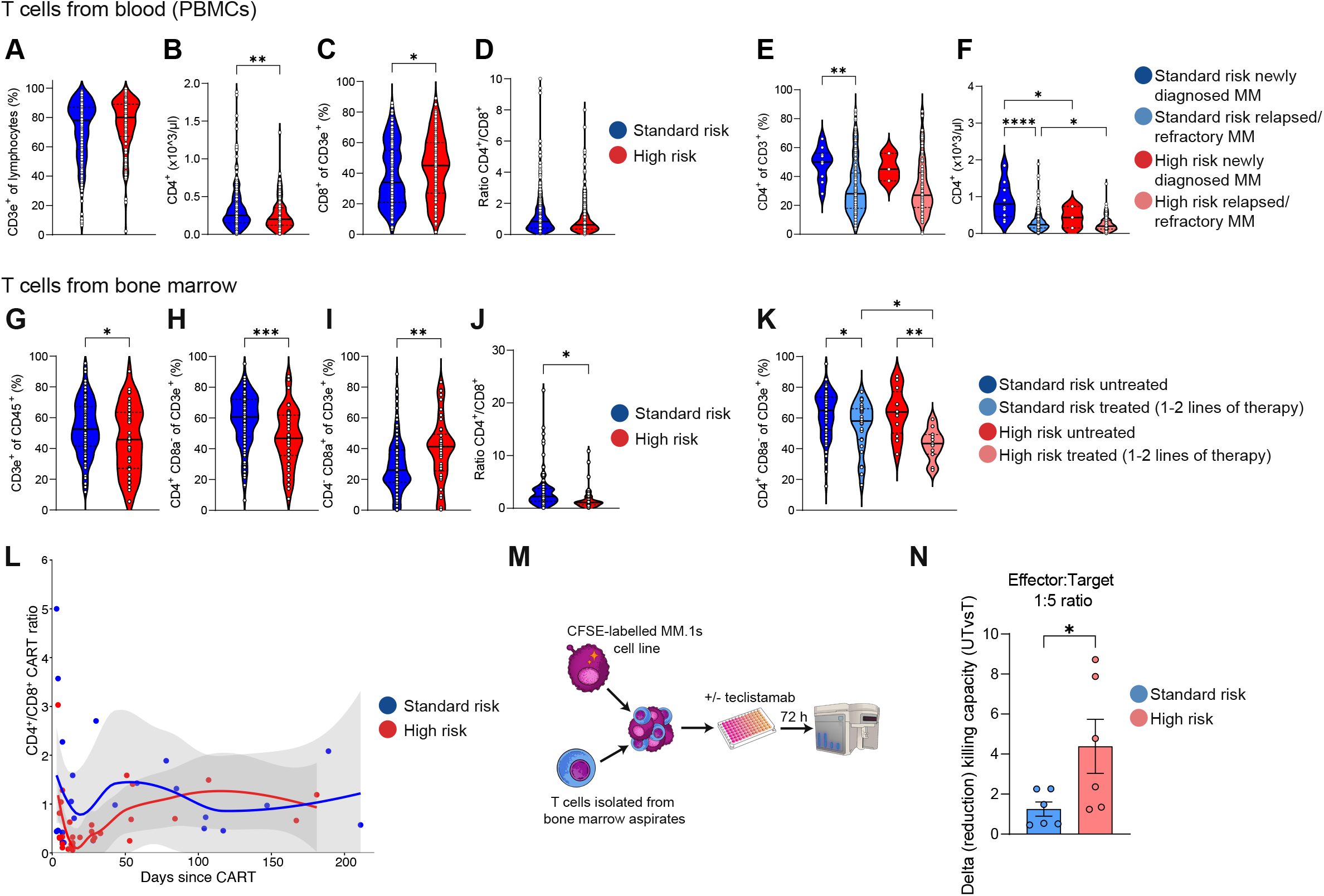
Loss of CD4+ T cells is more pronounced in high-risk patients. (**A** to **D**) Quantification of flow cytometry results from PB with patients separated by their risk status based on the SKY92 MMProfiler. Percentage of CD3+ T cells of lymphocytes (SR n=232; HR n=137) (A), absolute number of CD4+ T cells (SR n=207; HR n=115) (B), percentage of CD8+ of CD3+ (SR n=230; HR n=135) (C), and ratio CD4+ to CD8+ T cells (SR n=229; HR n=132) (D). Two-sided unpaired t-test was used for statistical testing. (**E** and **F**) Quantification of flow cytometry results from PB with patients being separated by their risk status based on the SKY92 profiler and NDMM (darker colors) and RRMM (lighter colors). Percentage of CD4+ of CD3+ (SR NDMM n=10; SR RRMM n=222; HR NDMM n= 3; HR RRMM n=133) (E) and absolute number of CD4+ T cells (SR NDMM n=10; SR RRMM n=197; HR NDMM n= 3; HR RRMM n=112) (F). Ordinary one-way ANOVA with Fisher’s LSD as a Post-hoc test was used for statistical testing. (**G** to **J**) Quantification of flow cytometry results from BM with patients separated by their risk status based on the SKY92 MMProfiler. Percentage of CD3e+ of CD45+ (SR n=117; HR n=45) (G), percentage of CD4+ of CD3e+ (SR n=117; HR n=46) (H), percentage of CD8a+ of CD3+ (SR n=117; HR n= 46) (I), and ratio CD4+ to CD8a+ T cells (SR n=116; HR n=45) (J). Two-sided unpaired t-test was used for statistical testing. (**K**) Flow cytometry quantification of percentage CD4+ of CD3e+ T cells from BM with patients separated by risk status and treatment status (untreated vs 1-2 lines of therapy) (SR untreated n=73; SR 1-2 lines of therapy n=28; HR untreated n=12; HR 1-2 lines of therapy n=11). Ordinary one-way ANOVA with Fisher’s LSD as a Post-hoc test was used for statistical testing. (**L**) Correlation between CD4+/CD8+ CAR-T cells and days after CAR-T infusion. Red color indicates high-risk patients, and blue color indicates standard-risk patients. Data was replotted from Xiang et al., 2024 *(17)*. (**M**) Experimental setup for *in vitro* cytotoxicity assay (killing assay). (**N**) Quantification of *in vitro* cytotoxicity assay by plotting delta (reduction) in killing capacity of T cells from treated MM patients compared to T cells isolated from untreated MM patients. Reduction was calculated by comparing killing capacity within the same risk group (n=6 per condition). Two-sided unpaired t-test was used for statistical testing. * p < 0.05, ** p < 0.01, *** p < 0.001, **** p < 0.0001.

In contrast, when analyzing the BM T cells, we observed a significant decrease in the CD3e+ T cell population in HR patients in comparison to SR patients (Fig. 2G). Furthermore, the decline in CD4+ T cells, the increase in CD8+ T cells, and the change in their ratio were all clearly evident (Fig. 2, H to J). We next compared patients that were ND (untreated) to patients that received 1-2 lines of therapy (treated) in the same cohort. We again found reduced proportions of CD4+ T cells (% of CD3e) in both SR and HR patients in the treated group compared to the untreated group. Significance was, however, also reached between SR and HR treated patients, indicating a more pronounced phenotype in HR patients (Fig. 2K). We further observed no differences in CD3+ absolute numbers, but an increase in CD8 T cell percentages (% of CD3e) and a decrease in the CD4+/CD8+ ratio in both SR and HR patients in the treated group compared to the untreated group (fig. S2, K to M). In summary, we observed a decline of CD4+ T cells in the PB and BM, which was associated with treatment and more pronounced in the BM and HR patients. Our findings thus suggest an altered BM immune microenvironment and circulating T cell compositions following anti-MM treatments especially in HR patients.

### A low CD4+/CD8+ T cell ratio in CAR-T cells is linked with high-risk disease and treatment impacts killing capacity *in vitro*

To determine if an decrease in the CD4+/CD8+ T cell ratio of the PB or BM can be translated into a decrease in the CD4+/CD8+ T cell ratio in the corresponding CAR-T cells, we plotted the ratio for SR and HR patients over the course of days following CAR-T cell infusion (Fig. 2L, *(17)*). In the first weeks after infusion, the CD4+/CD8+ CAR-T cell ratio was decreased in HR patients, but over time this ratio increased and was similar to the ratio found in SR patients. This indicates that CD4+ CAR-T cells are the more long-lived T cells *in vivo* in line with previous findings *(6)*. As changes in the CD4+/CD8+ T cell ratio might translate into reduced killing capacity due to less CD4 T cell help, we performed *in vitro* killing assays in the presence of the bispecific antibody teclistamab using mixed CD4+ and CD8+ T cells isolated from the BM of SR and HR patients untreated and treated. We labeled the myeloma cell line MM.1s with CFSE and co-cultured with the isolated T cells and teclistamab (Fig. 2M, fig. S2N). The killing capacity was reduced in patients that previously received treatment and this reduction was more prominent in HR patients seen in the larger delta killing capacity (Fig. 2N). In conclusion, the loss of CD4+ T cells resulted in unfavorable CD4+/CD8+ T cell ratios in CAR-T cells and reduced T cell-tumor cell killing capacity of isolated T cells in HR patients.

### ScRNAseq reveals profound differences in T cell subpopulations between standard- and high-risk patients before and after treatment

Given the high plasticity of T cell states, we used scRNAseq to perform a high-resolution analysis of their transcriptional landscapes in the BM. This allowed for the identification of discrete T cell states and programs. After fluorescence-activated cell sorting (FACS), equal numbers of live CD3e+ CD4+ and CD8+ T cells were collected from SR and HR patients receiving (treated) and not receiving (untreated) certain therapies (Fig. 3A, fig. S3A). All patients had a well-documented treatment history including their information of the SKY92 classifier (Fig. 3B). In total, we retained 36,277 T cells after quality filtering. Using the expression level of canonical marker genes, we obtained 19 T cell identities covering different CD8+ (clusters 10-18) and CD4+ (clusters 1-8) subsets, including CD4+ Tregs (cluster 9) *(18–22)* (Fig. 3C, fig. S3B). The cluster containing the most cells was cluster 8, CD4+ naïve T cells. Notably, we did not identify an exhausted T cell cluster as already shown by Pilcher et al. *(23)*. We then aimed to compare if there were changes in the proportion of clusters/subsets between untreated and treated patients within one risk group. Among SR patients, the CD4+ naïve T cell population decreased mildly, while all CD8+ effector memory CD45RA+ (EMRA; clusters 11-15) T cell populations increased markedly in treated patients compared to untreated patients. (Fig. 3D, left). Among HR patients, the CD4+ naïve T cell population decreased markedly, and the CD8+ naïve T cell population decreased mildly, while all CD8+ EMRA (clusters 11-15) T cell populations increased mildly in treated patients compared to untreated patients (Fig. 3D, right).

**Fig. 3.**
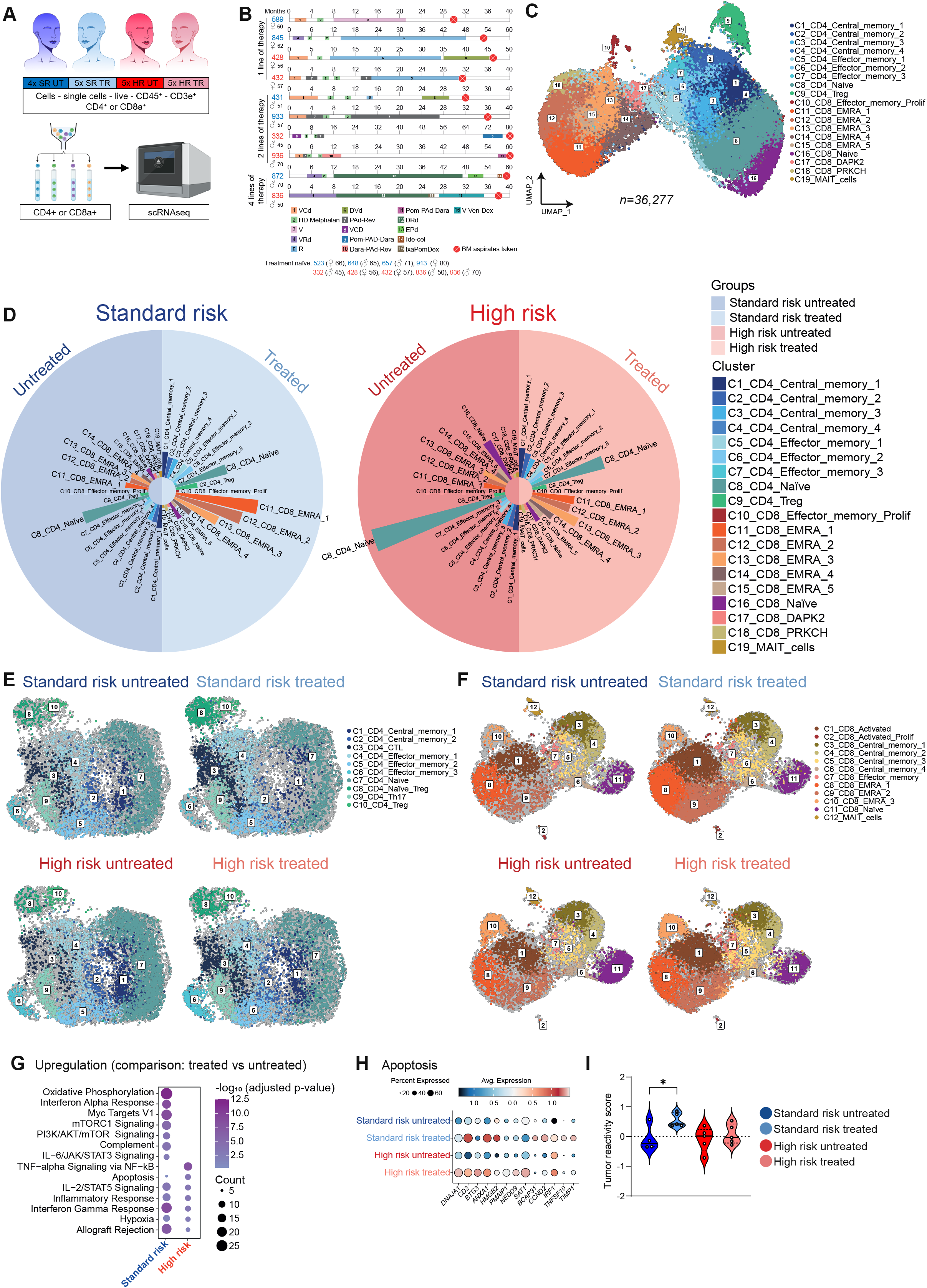
CD4+ naïve T cells upregulate apoptosis in treated high-risk patients. (**A**) Overview of experimental setup for scRNAseq. (**B**) Overview of treatments for each patient used for scRNAseq. (**C**) Uniform Manifold Approximation and Projection (UMAP) of all cells after quality control. (**D**) Circular barplot of clusters identified in (C). The left circular barplot represents standard-risk and the right circular barplot represents high-risk. Length of barplot indicates proportion of clusters compared to all clusters. (**E**) UMAP of separated CD4+ T cell clusters. (**F**) UMAP of separated CD8+ T cell clusters. (**G**) Pathway analysis with Hallmark gene sets of upregulated genes by comparing treated versus untreated patients within one risk group. (**H**) Differently expressed genes from “Apoptosis” pathway from (G). (**I**) Tumor reactivity score of CD8+ T cells based on gene signature from *(6)*. SR, standard-risk; UT, untreated; TR, treated; HR, high-risk; V, bortezomib; C, cyclophosphamide; d/Dex, dexamethasone; HD, high-dose; R, lenalidomide; D, doxorubicine; P, prednisone; A, AraC; Rev, revlimide; Pom, pomalidomide; Dara, daratumumab; E, etoposide; Ide-cel, Idecabtagen-Vicleucel; Ixa, ixazomib.

To get a deeper look into the two main T cell subsets, CD4+ or CD8+ T cells, we separated, re-clustered and re-annotated the populations (Fig. 3, E and F, fig. S3, C and G). Among CD4+ T cell subsets, CD4+ naïve T cells were reduced in treated SR and treated HR patients in comparison to untreated patients (Fig. 3E, fig. S3D). This loss of CD4+ naïve T cells was also validated with our flow cytometry data of the previously mentioned 162 MM patients (fig. S3, E and F, Fig. 1A). Among CD8+ T cell subsets, CD8+ naïve T cells were decreased in HR treated patients in comparison to untreated patients and CD8+ EMRA 1 and 2 T cells were increased in SR treated patients in comparison to untreated patients (Fig. 3F, fig. S3H). This was further supported by flow cytometry analysis (fig. S3I).

To examine transcriptional differences, we compared naïve CD4+ T cells from treated and untreated patients within each risk group and performed pathway enrichment analysis on the differentially upregulated genes using the Hallmark gene sets. Whereas SR treated CD4+ naïve T cells upregulated genes linked with oxidative phosphorylation and allograft rejection in comparison to those of untreated (Fig. 3G), indicating a fitter, more activated, T cell state, CD4+ naïve T cells from treated HR patients upregulated genes linked with apoptosis in comparison to those of untreated (Fig. 3, G and H). As we saw differences in *ex vivo* tumor cell killing (Fig. 2N), we harnessed our scRNAseq data as well to calculate a tumor reactivity score *(6)*. Interestingly, treatment did not have a negative impact on the tumor reactivity in both risk groups. However, only CD8+ T cells of SR treated patients in comparison to untreated patients increased their tumor reactivity score, while there was no difference in CD8+ T cells of HR patients (Fig. 3I). In summary, we found a striking loss of naïve CD4+ T cells in HR patients after treatment linked with induction of apoptosis programs. In contrast, in SR patients CD8+ EMRA T cells increased after treatment and showed higher tumor reactivity.

### Highly proliferating MM cell line upregulates cytokine and MHC II expression upon melphalan treatment and drive T cell inactivation

As melphalan showed the most pronounced effect on CD4+ T cell populations, we wanted to investigate the underlying mechanism. First, we investigated the direct effect of melphalan on MM cells. For this purpose, we used two MM cell lines: one relatively slow-growing cell line (MM.1s) and one fast-growing cell line (RPMI-8226, more indicative of HR disease; fig. S4, A to C). We performed inhibitory concentration 50 (IC50) viability curve analysis to determine the half-lethal concentration of melphalan on cell viability. As expected, higher concentrations were needed to reduce viability of the RPMI-8226 cell line, than the slow growing MM.1s cell line leading also to differences in IC50 values (Fig. 4A). We then used the IC25 to determine the transcriptional programs via bulk RNAseq. The two cell lines produced different profiles, with the most variance between them (Fig. 4, B and C). Surprisingly, melphalan treatment induced only marginal changes (9 upregulated differential expressed genes (DEGs) and 0 downregulated DEGs) in the MM.1s cell line. In the RPMI-8226 cell line, however, melphalan induced more drastic changes (296 upregulated [left] and 251 downregulated DEGs [right], Fig. 4D). The upregulated DEGs in RPMI-8226 after melphalan treatment were linked with pathways associated with mostly inflammatory responses (Fig. 4E). Looking further into the genes involved in the top pathways, we found that they were either cytokines and chemokines or MHC II associated molecules indicating enhanced cross-talk to CD4+ T cells (Fig. 4F).

**Fig. 4.**
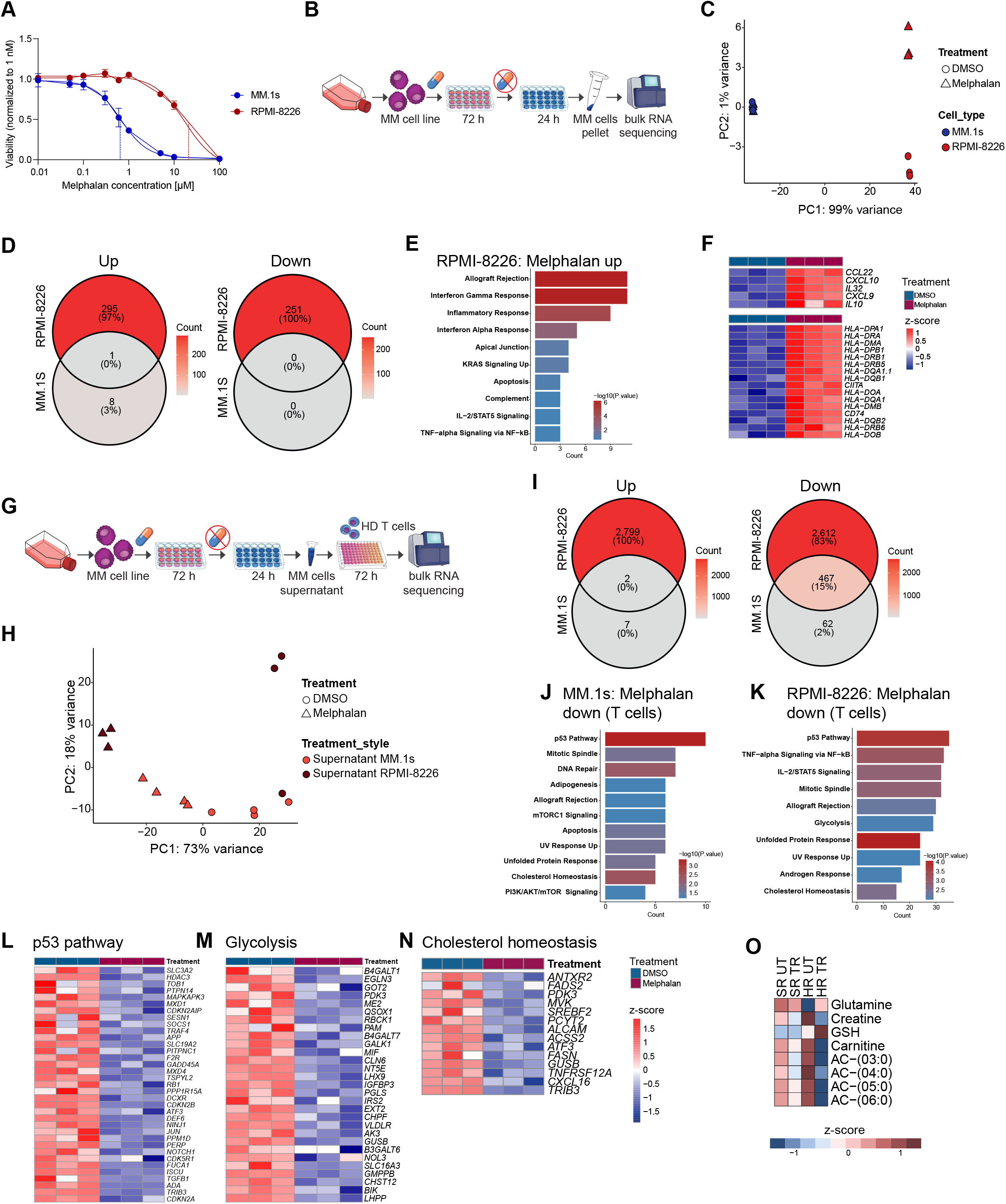
RPMI-8226 upregulates cytokine and MHC II expression upon melphalan treatment and drives T cell inactivation. (**A**) Viability of MM cell lines treated with different melphalan concentrations. MM.1s cell line in blue and RPMI-8226 in red. Dotted line indicates IC50 concentration. **(B)** Bulk RNA sequencing setup for MM cell lines (MM.1s and RPMI-8226) treated with melphalan. (**C**) Principal component analysis (PCA) of bulk RNA sequencing of MM cell lines after DMSO or melphalan treatment. Blue indicates MM.1s and red indicates RPMI-8226 cell line. Points indicate DMSO treatment and triangles indicate melphalan treatment (n=3-4 per condition). **(D)** Venn diagram from up- and downregulated genes in MM cell lines (comparing melphalan to DMSO). **(E)** Hallmark gene sets analysis of upregulated genes in melphalan treatment from RPMI-8226. **(F)** Heatmap of genes linked with inflammatory response pathway and MHC II-related genes upregulated in melphalan treatment. **(G)** Bulk RNA sequencing setup for T cells treated with the supernatant of melphalan or DMSO treated MM cell lines. **(H)** PCA plot of sequenced T cells. Light red indicates treatment of T cells with supernatant of treated MM.1s and dark red indicates treatment of T cells with supernatant of treated RPMI-8226. Points indicate DMSO treatment and triangles indicate melphalan treatment (n=3-4 per condition). **(I)** Venn diagram of up- and downregulated genes from supernatant treated T cells. **(J)** Hallmark gene sets analysis of downregulated genes from with MM.1s supernatant treated T cells. **(K)** Hallmark gene sets analysis of downregulated genes from with RPMI-8226 supernatant treated T cells. **(L)** Heatmap of genes linked with p53 pathway downregulated in melphalan treatment (K). **(M)** Heatmap of genes linked with glycolysis pathway downregulated in melphalan treatment (K). **(N)** Heatmap of genes linked with cholesterol homeostasis pathway downregulated in melphalan treatment (K). **(O)** Heatmap of z-scores from water-soluble metabolites from the BM plasma of SR and HR patients of selected significant metabolites.

In the next step, we wanted to examine this cross-talk more closely and thus transferred the supernatant of both MM cell lines after melphalan treatment and a wash-out period of 24h onto PB healthy donor T cells (Fig. 4, G and H). The effect of the supernatant from MM.1s had less effect on T cells than the supernatant from RPMI-8226 depicted again by the number of found DEGs. Overall, more DEGs were found to be downregulated after the treatment with the supernatant of RPMI-8226 treated with melphalan in comparison to treated with DMSO (Fig. 4I). The associated downregulated pathways of the supernatant-treated T cells were similar between both cell lines, even though the RPMI-8226 cell line led to a higher gene count per pathway. These indicated major defects in cell cycle progression (pathways p53 and mitotic spindle) and metabolism (pathways glycolysis and cholesterol homeostasis; Fig. 4, J to N). Glycolysis is a main metabolic pathway for T cell function and proliferation *(24, 25)* (Fig. 4M). Furthermore, cholesterol homeostasis plays an important role in naïve T cell metabolism upon T cell activation *(26)* (Fig. 4N). As many metabolism related pathways were found downregulated in T cells after supernatant treatment of RPMI-8226 melphalan treated cells, we compared water-soluble metabolites from the BM plasma between SR and HR patients in the treated and untreated setting. Indeed, we found plasma of HR patients after treatment to be associated with a depletion of certain lipid-associated metabolites, such as carnitine and several acyl-carnitines, while redox important factors, such as glutathione (GSH), were increased, but we also found enrichment (arginine, histidine, and lysine) or depletion (glutamine) of certain amino acids (Fig. 4O, full list fig. S4D). Together, our data indicate a more severe stress for T cells in a HR-setting treated with melphalan. Furthermore, we demonstrate that melphalan induces programs of cross-talk with CD4+ T cells in high-proliferating PCs with induced secreted and metabolic factors negatively impacting T cell function.

### Induction of redox stress induces loss of naïve CD4+ T cells

To investigate if HR disease is as well associated with an adapted nutrient environment, changed lipids and redox factors, we re-analyzed a publicly available micro-array dataset from BM PCs, including risk classification into HR or SR. We then calculated a SKY92 score based on the 92 genes from the SKY92 MMProfiler. As expected, the score was enriched in the PCs of HR patients (Fig. 5A). The PCs from HR patients upregulated genes linked with pathways that were associated with proliferation, such as E2F and Myc targets, but also oxidative phosphorylation, glycolysis, fatty acid metabolism, and ROS pathway in HR PCs, suggesting a fast growing capacity which inherently is associated with depletion of T cell necessary nutrients and enrichment in ROS production (Fig. 5, B and C, fig. S5A). We next wanted to quantify this ROS production in our fast-growing MM cell line RPMI-8226 (more HR disease-like) using the CellROX dye. Indeed, ROS levels were evident and in fact further increased after melphalan treatment (Fig. 5D). Furthermore, the effect on viability of melphalan on the MM cell lines RPMI-8226 and MM.1s could be reduced by the addition of the antioxidant N-acetyl-L-cysteine (NAC), resulting in increased IC50 values (fig. S5B).

**Fig. 5.**
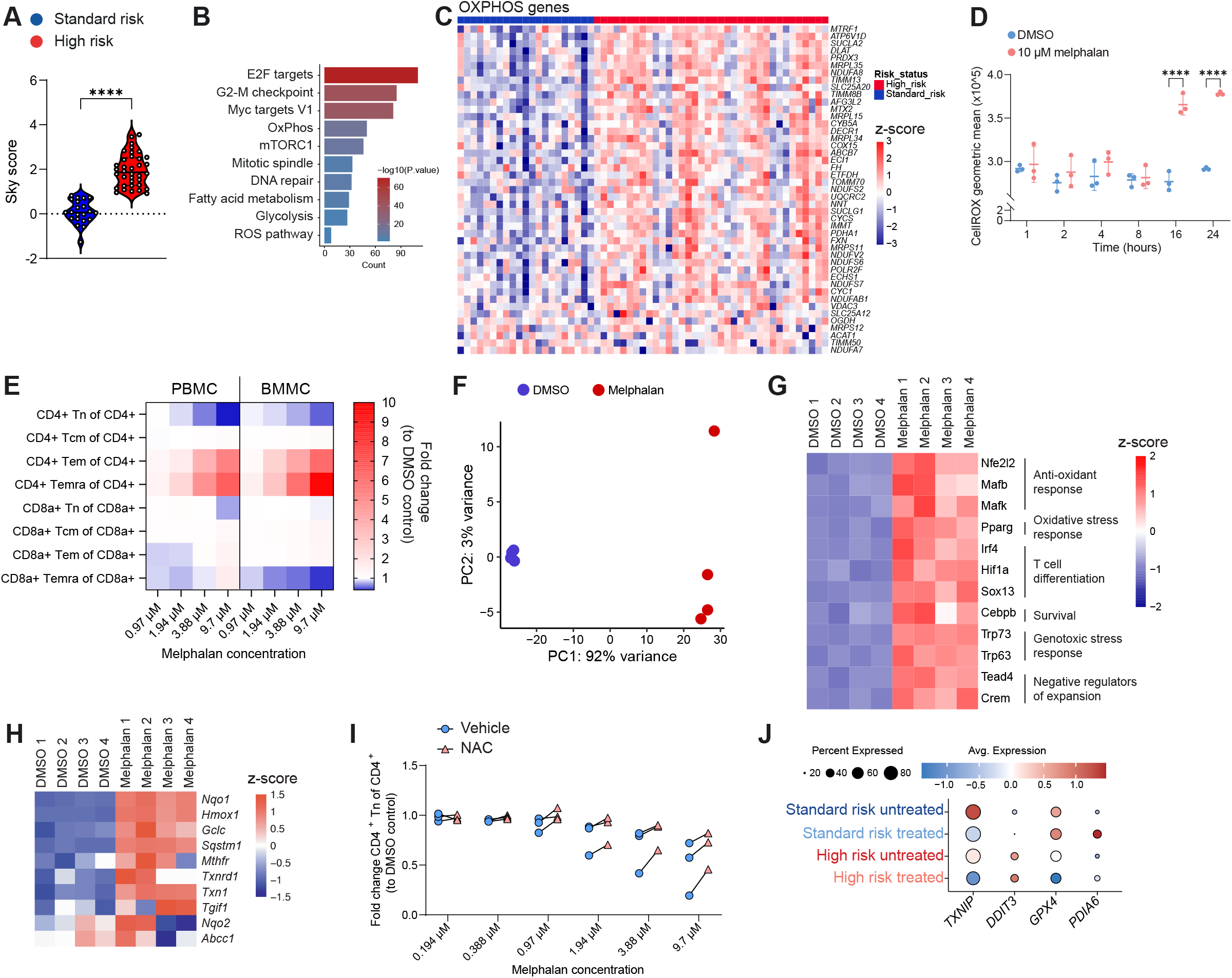
Redox stress induction leads to depletion of CD4+ naïve T cells but can be attenuated by supplementation of N-acetyl-L-cysteine. **(A)** Violin plot of SKY92 gene signature *(16)* comparing HR to SR disease in GSE164703 microarray data set *(55)*. Two-sided unpaired t-test was used for statistical testing. **(B)** Hallmark gene sets analysis of upregulated genes in PCs in HR patients from GSE164703 microarray data set *(55)*. **(C)** Heatmap of genes linked with oxidative phosphorylation pathway from (B). **(D)** CellROX staining of RPMI-8226 treated with 10 µM melphalan or DMSO. Treatment was performed for indicated time periods (hour, n=3 for each timepoint). Two-way ANOVA with Fisher’s LSD as a Post-hoc test was used for statistical testing. (**E**) Heatmap representing the fold change in the proportion of T cells compared to vehicle control following the *in vitro* treatment of PB and BM mononuclear cells with melphalan (n=3 for each group). (**F**) PCA of murine CD4+ naïve T cells treated with 10 µM melphalan or DMSO. Blue indicates DMSO treatment, red indicates melphalan treatment (n=4 for each condition). **(G)** Heatmap showing top most variable transcription factor (TF) activities across the samples. **(H)** Heatmap showing differential target genes of Nfe2l2 TF. **(I)** *In vitro* treatment of PB mononuclear cells with melphalan and 1 mM N-acetyl-L-cysteine (NAC). Blue circles indicate melphalan treatment, orange triangles indicate melphalan and NAC treatment (n= 3 replicates per condition). **(J)** Average expression of specific genes from scRNAseq of sorted T cells from BM. **** p < 0.0001.

To clearly delineate why CD4 and in particular naïve CD4 T cells were most sensitive to treatment both in case of HR disease and melphalan, we also tested the impact of melphalan on CD4+ and CD8+ T cells and their subsets directly. T cells isolated from the PB of three healthy donors or isolated from the BM of three NDMM patients were treated with six increasing concentrations of melphalan (Fig. 5E). The used concentrations of melphalan had no effect on the viability of T cells (fig. S5C). Naïve CD4+ T cells declined, while effector memory (EM) and effector memory naïve-like (EMRA) CD4+ T cells with increasing concentrations of melphalan in both PB and BM T cells increased (Fig. 5E and fig. S5D). On the contrary, naïve CD8a+ T cells were less sensitive (PB) or not at all (BM), while exclusively EMRA CD8+ T cells of the BM declined with increasing melphalan concentrations (Fig. 5E and fig. S5D). To investigate the transcriptomic changes induced by melphalan in naïve CD4+ T cells specifically, we isolated these cells from murine spleens, treated with 10 µM melphalan for three days and performed bulk RNAseq (Fig. 5F). We found in total 2804 DEGs (with abs log2 fold change greater than 1; p < 0.05), of which 1744 were upregulated and 1060 were downregulated. To infer changes in transcription factor (TF) activity, we then performed a TF enrichment analysis based on the expression profile of the bulk RNAseq data and subsequently identified the most variable TFs across conditions (fig. S5E). We found many TFs that were either associated with oxidative stress or T cell differentiation/effector function (Fig. 5G). In particular, within the signature of Nfe2l2 (NRF2), the major TF driving cellular defense against toxic and oxidative insults, we found genes associated with the antioxidant response, such as *Nqo1, Txnrd1* and *Txn1* (Fig. 5H). This suggested that oxidative stress might be a reason for the loss of CD4+ naïve T cells after melphalan treatment. We next wanted to test whether reduction of ROS would also rescue melphalan induced naïve CD4+ T cell reduction and used again NAC treatment. Interestingly, ROS scavenging via NAC indeed rescued the naïve CD4+ T cell loss induced by melphalan (Fig. 5I).

Together, the data suggested a combinatorial effect of HR PCs and melphalan on naïve CD4+ T cells. In fact, this was supported by enrichment of extracellular glutathione (GSH) in the BM plasma of HR patients, supporting a higher redox stress in the BM plasma of HR patients after treatment (Fig. 4O). We went back to our scRNAseq data of BM T cells of patients with different risk status and treatment schedule. Interestingly, *TXNIP*, an inhibitor of thioredoxin, was highly downregulated in T cells of HR patients after treatment, indicating an active antioxidant response by thioredoxin in these T cells. Furthermore, *DDIT3*, a gene linked with endoplasmic reticulum stress and apoptosis, was upregulated in HR patients’ T cells. While *GPX4* and *PDIA6* were highly expressed in SR patients’ T cells indicating less lipid peroxidation and improved development of these T cells *(26, 27)*(Fig. 5J).

In summary, naïve CD4+ T cells appear to be the most sensitive to melphalan among all T cell subsets due to the induction of an oxidative stress response, which is most pronounced in HR disease. The reason for this is that HR PCs inherently produce more ROS and deplete nutrients due to increased proliferation. Therefore, an overexaggerated ROS and T cell unfavorable environment may lead to the apoptosis of naïve CD4+ T cells specifically in HR disease after melphalan treatment.

## DISCUSSION

In this study, we show the negative impact of prior treatment on CD4+ T cells, in particular naïve CD4+ T cells, in the PB and BM of MM patients. The reduction in this population was more pronounced in melphalan-treated patients and/or patients with HR disease. However, T cell based therapies, such as bsAbs or CAR-T cells, heavily rely on functional endogenous T cells.

Specifically, it has recently been shown that a high CD4+ T cell count is critical for CAR-T cell production *(13)*, as well as tumor killing and durable disease control *(6)*. Moreover, CD4+ T cells exhibit superior longevity *in vivo* compared to other CAR-T subsets *(14)* and compared to CD8+ CAR-T cells do not become apoptotic or exhausted *(28)*. In contrast, high numbers of CD4+ T cells in CAR-T cell products have been linked to immune-related adverse events associated with CAR-T cell therapy, suggesting that an adequately balanced amount of CD4+ T cells is critical to achieve a durable response without over-stimulation *(29)*.

Melphalan is the cornerstone of conditioning for autologous stem cell transplantation (ASCT), which is standard of care for fit newly diagnosed patients *(30–34)*. High-dose melphalan provides a level of disease eradication and depth of response that remains difficult to replicate with other MM therapies. However, the use of high-dose melphalan has been called into question in the era of modern therapies *(35)*. Thus, rather than excluding melphalan from the treatment, alternative options include CAR-T cell production or leukapheresis prior to high-dose chemotherapy treatment, at least for patients with HR disease.

ROS are key to T cell signaling, as following T cell receptor (TCR) stimulation, T cells increase ROS production with mitochondria serving as the central oxidative signaling platform. This ROS then signals to REDOX-sensitive transcription factors, such as NFAT, NF-κB and AP-1 necessary for T cell activation *(36)*. Long-term exposure of CD4+ T cells to ROS results in the selective inhibition of the DNA-binding ability of NFAT and NF-κB, which leads to the downregulation of IL-2 transcription *(37)*. This indicates that ROS levels play a pivotal role in CD4+ T cell activation and differentiation, and that this balance needs to be fine-tuned.

In our study, metabolites critical for T cell function were depleted in untreated HR patients, a phenomenon no longer observed in treated HR patients. This suggested a significant divergence from the SR patients. Interestingly, it was shown that protein starvation is associated with infections and a diminished vaccine response *(38, 39)*. The administration of protein supplements has been identified as a potential strategy to enhance this outcome *(40)*. In general, therapy induced oxidative stress in our patients, which could indicate that supplementation with a substance such as creatine or NAC could rescue this by exerting an antioxidant effect *(41)*.

The decrease of CD4+ T cells, including naïve CD4+ T cells, in the PB of patients undergoing chemotherapy has been previously described but the underlying mechanism remained elusive *(7, 42)*. In B-cell lymphoma, this decline was largely attributed to mitochondrial damage, as shown by reduced oxygen consumption induced by cyclophosphamide, doxorubicin or cytarabine *(43)*. Here, we attribute this loss in both the PB and BM to melphalan, and possibly also to carfilzomib.

In conclusion, alterations in the T cell composition were identified, with a loss of CD4 naïve T cells in a treatment-dependent manner in HR disease. This might be attributable to the metabolite tumor microenvironment, which provides a rationale for collecting T cells before treatment with melphalan to manufacture an optimal CAR-T cell product.

## MATERIALS AND METHODS

### Study design

Peripheral blood and bone marrow samples were collected following the Declaration of Helsinki and approved by the review board of the University of Würzburg (Germany, reference no. #197/20-am). All patients gave their written informed consent.

Peripheral blood samples were freshly analyzed in the immune diagnostics of the University Hospital Würzburg. Bone marrow samples were used for risk stratification using the SKY92 MMProfiler and negative fractions, which did not contain CD138+ plasma cells, were frozen and stored at −150°C. To analyze the T cell populations, samples from the peripheral blood and bone marrow were analyzed with flow cytometry. Cytotoxic functions of T cells from the bone marrow were used in cytotoxic assays (n=20). Single cell RNA sequencing was performed on age- and gender-matched patients, who were either untreated or received up to four lines of therapy (n=20). To study the effect of melphalan on myeloma cell lines and T cells isolated from healthy donor peripheral blood, myeloma cell lines (n=2) and healthy donor T cells were treated with an inhibitory concentration 25 (IC25), which was calculated from drug screen experiments. After treatment RNA was isolated and then further processed for bulk RNA sequencing. Water-soluble metabolites were measured with liquid chromatography-mass spectrometry to identify enriched or depleted metabolites in the bone marrow plasma (n=49). To validate results from scRNAseq, peripheral blood and bone marrow mononuclear cells were treated *in vitro* with different concentrations of melphalan. CD4+ naïve T cells were isolated from a murine spleen and treated with melphalan to examine the specific effect on this subpopulation. Melphalan is known to induce redox stress, therefore CellROX stainings were performed on one myeloma cell line. The redox stress induced CD4+ naïve T cell depletion was rescued by addition of N-acetylcysteine.

### Statistical analysis

Statistical analysis was performed with GraphPad Prism version 9 (Insight Partners) or R v4.5.2 *(60)*. Used tests include: two-sided unpaired t-test, ordinary one-way ANOVA, ordinary two-way ANOVA, or log-rank test. Tests used for each figure are indicated. Following abbreviations were used for p-values: * p < 0.05, ** p < 0.01, *** p < 0.001, **** p < 0.0001.

### List of Supplementary Materials

Figs. S1 to S5

Tables S1 to S3

## Acknowledgments

*We warmly thank the patients who participated in this study. MJ, NA, MJK, MH, KMK, LR, and AR thank the German Cancer Aid via the MSNZ program. LR thanks the Else Kröner Forschungskolleg Twinsight for support. In addition, we thank the FACS Core Unit of the IZKF Würzburg for the support of this study. This study was funded by the IZKF-Project Z-12. We further acknowledge the sequencing efforts of the Core Unit Systems Medicine (IZKF Z-6) and especially Panagiota Arampatzi. We also thank the immune diagnostic lab of the polyclinic of the University Hospital Würzburg*.

## Author contributions

Conceptualization: MJ, LR, AR

Methodology: MJ, NA, XZ, ES, HF, AA

Formal analysis: MJ, NA, XZ, MJK, MHe

Investigation: MJ, XZ, ES, HF, WS, LG, MHe, AA, AW, SW, BME, AH, MS

Visualization: MJ, NA, AR

Funding acquisition: XZ, HE, LR, AR

Project administration: MJ, XZ, LR, AR

Resources: XZ, ES, WS, SN, CV, SK, MJS, MHu, HE, KMK, LR

Supervision: LR, AR

Writing – original draft: MJ, NA, XZ, LR, AR

Writing – review & editing: all authors contributed

## Data and materials availability

All data associated with this study are present in the paper and the supplementary materials. All sequencing data will be available in the Genome Sequencing Archive (GSA) upon acceptance of the paper. All materials used or generated in this study are commercially available or will be supplied upon reasonable request.

