## Supplementary files for "Profound CD4+ T-Cell Reprogramming by Melphalan-Driven Oxidative Stress in High-Risk Multiple Myeloma"

### Supplementary materials:

Material and methods

Figs. S1 to S5

Tables S1 to S3

References (*44-60*)

**Material and methods:**

**BMMC isolation and SKY92 profiling**

Bone marrow aspirates were spun down at 1000 rcf for 10 min. 2 ml of plasma were immediately frozen at -80°C. The remaining sample was mixed 1:1 with DPBS (ThermoFisher, Cat. No. 14190169), filtered (100 µm filter, Greiner, Cat. No. 542000) and added on 15 ml density gradient medium (Anprotec, Cat. No. AC-AF-0018). Samples were spun down at 800 rcf for 20 min with break off. Mononuclear cell layer was transferred into a new tube and washed twice with DPBS. BMMCs were diluted to 1 x10^8 cells/ml. CD138 positive plasma cells were isolated with the Human CD138 Positive Selection Kit II (Stemcell, Cat. No. 17877) with the RoboSep (Stemcell, Cat. No. 38008). After isolation, CD138+ cells were used for risk status determination and negative fraction cells were frozen in CryoStor cell cryopreservation media (Sigma-Aldrich, Cat. No. C2874). For risk stratification, RNA from CD138 positive cells was isolated with the QIAGEN AllPrep DNA/RNA Micro Kit (QIAGEN, Cat. No. 80284) and was used with the MMprofiler CE IVD Assay Kit (SkylineDx BV, Cat. No. IFU-012) following manufacturer's protocol.

**Flow cytometry**

Negative fractions (BMMCs) were thawed at 37°C and transferred to 9 ml pre-warmed MM medium (RPMI-1620 (ThermoFisher, Cat. No. 11875093) + 10% FBS (Sigma-Aldrich, Cat. No. F7524) + 1% Penicillin/Streptomycin (ThermoFisher, Cat. No. 15140122) + 1% Sodium pyruvate (ThermoFisher, Cat. No. 11360070) + 1% GlutaMax (ThermoFisher, Cat. No. 35050061)). Cells were spun down at 400 rcf for 10 min at RT. Supernatant was removed and pellet was resuspended in 1 ml MM medium with 50 µg DNase I to achieve single cell suspension. Digestion was incubated for 5 min at 37°C while shaking and stopped by adding 5 mM EDTA and then 9 ml pre-warmed MM medium was added. Cells were spun down with the same settings. Supernatant was removed and pellet was resuspended in 400 µl MM medium. Cell suspension was equally distributed on 2x 96 well suspension plates. Plates were spun down at 532 rcf for 2 min at 4°C. Supernatant was removed, pellet was resuspended in 100 µl DPBS (Sigma-Aldrich, Cat. No. D8537) + live/dead stain (Fixable viability die, APC-eFluor780 ThermoFisher, 1:1000, Cat. No. 65-0865-14) + Human TruStain FcX™ (Biolegend, 1:20, Cat. No. 422302) and incubated for 15 min at 4°C. Cells were washed once with DPBS and the antibody mix in FACS buffer (DPBS + 2 mM EDTA (Merck, Cat. No. 324504) + 0.5% BSA (Roche, Cat. No. 10735086001)) was added ([α-human-CD3ε, Alexa Fluor (AF) 700, clone: HIT3a, 1:300, Biolegend, Cat. No. 300324], [α-human-CD4, Brilliant Violet (BV) 421, clone: Okt4, 1:300, Biolegend, Cat. No. 317434], [α-human-CD8a, BV605, clone: SK1, 1:300, Biolegend, Cat. No. 344742], [α-human-CD45, PE, clone: 2D1, 1:100, Biolegend, Cat. No. 368510], [α-human-CD45RA, eFluor506, clone: HI100, 1:200, ThermoFisher, Cat. No. 69-0458-41], [α-human-CD62L, SB600, clone: DREG-56, 1:100, ThermoFisher, Cat. No. 63-

0629-41], [α-human-CD8a, BV650, clone: RPA-T8, 1:20, Biolegend, Cat. No. 301042]). Antibodies were stained for 30 min at 4°C. Cells were washed once with FACS buffer and then measured at the Attune™ NxT Flow Cytometer (ThermoFisher, Cat. No. A29004). Flow cytometry data was analyzed with FlowJo v10.8.1 (BD) and visualized with GraphPad Prism 9 (Insight Partners).

For peripheral blood samples, 100 µl EDTA blood was used in DuraClone IM Phenotyping BASIC tube (Beckman Coulter, Cat. No. B53309) (containing: [α-human-CD45, Krome orange dye, clone: J33, Beckman Coulter], [α-human-CD3, APC-A700, clone: UCHT1, Beckman Coulter], [α-human-CD8, Pacific blue, clone: B9.11, Beckman coulter], [α-human-CD4, APC, clone: 13B8.2, Beckman Coulter]) following manufacturer's protocol.

**Progression-free survival, overall survival and multivariate testing**

Kaplan-Meier curves for progression-free survival and overall survival were generated using the ggsurvplot2 function from survtools 0.1 R package *(44)*. Multivariable associations were assessed using a generalized linear model (GLM) with a Gamma distribution and log link and implemented by stats 4.5.2 R package *(45)*. Effect estimates were exponentiated and reported as expected mean ratios (EMRs) with 95% confidence intervals. Forest plot was generated using the forestploter 1.1.4 R package *(46)*.

**Killing assays**

MM.1s cells were collected and counted. Cells were spun down at 400 rcf for 5 min at RT and then resuspended in pre-warmed DPBS for a final concentration of 1 Mio cells/ml. CellTrace™ CFSE (1:1000, ThermoFisher, Cat. No. C34554) was added and incubated for 20 min at 37°C in the dark. Pre-warmed MM medium was added (5x volume of staining solution) and incubated for 5 min at RT. Cells were spun down (400 rcf, 5 min, RT) and resuspended in pre-warmed MM medium to a final concentration of 1 Mio cells/ml. Cells were seeded depending on effector-to-target (E:T) ratio (1:5, 1:10) on a 96 well plate. Negative fractions were thawed as described for flow cytometry. After thawing, samples were then processed with the human Pan T cell isolation kit following manufacturer's protocol (Miltenyi, Cat. No. 130-096-535). Isolated T cells were then resuspended in pre-warmed MM medium to a final concentration of 1 Mio cells/ml and seeded according to the E:T ratio. 1 µM Teclistamab was added and DPBS was used as negative control. Reactions were incubated for 3 days. Plate was spun down at 532 rcf for 2 min at 4°C and supernatant was removed. Cells were washed once with DPBS and then stained with live/dead stain (Fixable viability die, APC-eFluor780 ThermoFisher, 1:1000) in DBPS for 15 min at 4°C. Afterwards, cells were washed once with FACS buffer and resuspended in FACS buffer and measured at the Attune™ NxT Flow Cytometer. Flow cytometry data was analyzed with FlowJo v10.8.1 (BD) and visualized with GraphPad Prism 9 (Insight Partners). Specific lysis was calculated by following equation: Specific lysis (%) = [1 – (MM cell numberTeclistamab/MM cell numberDPBS)] x 100. Specific lysis was further normalized by CD8a+ T cells proportion detected with flow cytometry following this equation: Specific lysis (%)/CD8a+ of CD3e+ T cells (%).

**scRNAseq**

Cells were thawed as described in flow cytometry. Cells were counted after digestion and then spun down at 400 rcf for 10 min at RT. Pellets were resuspended in 100 µl live/dead solution (Fixable viability dye, APC-eFluor780, 1:1000) and incubated for 15 min at 4°C. Cells were spun down 400 rcf for 5 min at 4°C. Cells were washed once with DPBS + 10% FBS.

Afterwards, cells were blocked with Human TruStain FcX™ (1:20 in DPBS) for 10 min at 4°C. Antibody solution (containing antibodies ([α-human-CD3ε, FITC, clone: HIT3a, 1:300, Biolegend, Cat. No. 300306], [α-human-CD4, BV421, clone: Okt4, 1:300, Biolegend, Cat. No. 317434], [α-human-CD8a, APC, clone: RPA-T8, APC, 1:300, ThermoFisher, Cat. No. 17-0088-42], [α-human-CD45, PE, clone: 2D1, 1:300, Biolegend, Cat. No. 368510]) and hashing antibodies [TotalSeq™-B0251 anti-human Hashtag 1 Antibody, Biolegend, Cat. No. 394631], [TotalSeq™-B0252 anti-human Hashtag 2 Antibody, Biolegend, Cat. No. 394633], [TotalSeq™-B0253 anti-human Hashtag 3 Antibody, Biolegend, Cat. No. 394635], [TotalSeq™-B0254 anti-human Hashtag 4 Antibody, Biolegend, Cat. No. 394637]) was added to blocking solution and incubated for 30 min at 4°C. Cells were washed again with DPBS + 10% FBS and then resuspended in DPBS + 2% BSA. Before sorting, cells were filtered (40 µm, pluriSelect, Cat. No. 43-57040-01). 10,000 cells per population and patient were sorted at the FACSAria™ III Cell Sorter (BD) (Alive -> CD45 positive -> CD3ε positive -> CD8a or CD4 positive). After sorting, cells were counted and 20,000 cells (pooled from 4 patients) were loaded on the Chromium chip, following the manufacturer's protocol (10X Genomics, CG000317 Rev D). Quality of the library was checked with Bioanalyzer High Sensitivity DNA Kit (Agilent, Cat. No. 5067-4626). Libraries were sequenced on the NextSeq2000 (Illumina, San Diego, USA) with paired-end sequencing (26 bp and 90 bp).

**scRNAseq analysis**

Following sequencing, cell ranger 7.0.0 was used to map the libraries, using GRCh38 (GENCODE v32/Ensembl 98) as a reference. The workflow from the Seurat 5.1.0 R package was used for the analysis of the data *(47)*. Cells with fewer than 300 and more than 5000 genes, as well as those with more than 7.5% mitochondrial genes, were eliminated for quality control. To summarize, the data were log-transformed and normalized by a scale factor of 10,000. Then, "vst" was used to identify the top 2000 variable features. For the doublets detection and demultiplex cells to their origin, we applied HTODemux function from Seurat. Principal component analysis (PCA) was performed, and the top 50 principal components were selected for the further analysis. UMAP embedding for the dimensional reduction was applied on these top 50 principle components. To integrate cells by removing batch effects from patients, the Harmony algorithm was applied. Next, based on the expression of the top marker genes in each cell cluster found by FindMarkers function from Seurat package, all the cell clusters were annotated with different cell types. Lastly, the top upregulated genes with a log2 fold change cutoff 0.5, which are expressed in at least 30% cells comparing the treated samples with untreated samples in each of the risk status were considered to perform pathway enrichment analysis using clusterProfiler's v4.16.0 enrichment tools *(48)*. To evaluate the enrichment of a gene signature per individual cell, we computed a score using the AddModuleScore function from the Seurat R package. Lastly, ggplot2 and SCpubr were utilized for the visualization of the data *(49, 50)*.

**Doubling experiment**

100,000 cells of RPMI-8226 and MM.1s were seeded in 1 ml MM medium and were counted every 24 hours for 72 hours by manual counting.

**Bulk RNA assay setup myeloma cell lines and healthy donor T cells**

3 healthy donor PBMCs were cultivated for 14 days in T cell medium (RPMI-1620 + 10% human plasma + 1% Penicillin/Streptomycin) + 1% Sodium pyruvate + 1% GlutaMax + 150

IU/ml IL-2) in a α-CD28 (1:200 in DPBS, ThermoFisher, Cat. No. 16-0289-85) coated 96 well plate. Medium was changed every two days. After 14 days, cells were frozen in 90% FBS + 10% DMSO. For bulk RNAseq experiment, cells were thawed in 9 ml RPMI-8226 and spun down at 400 rcf for 5 min at 4°C. Supernatant was removed and pellet was resuspended in 1 ml PlasmaX™ (CancerTools, Cat. No. 156371) + 10% human plasma (LGC Seracare, Cat. No. 1810-0013) + 1% Penicillin/Streptomycin and counted. Cells recovered overnight at 37°C at a concentration of 2 Mio cells/ml. The next day, cells were spun down at 400 rcf for 5 min at 4°C. Supernatant was removed and pellet was resuspended in 2 ml RPMI-8226 with 50 µg DNase I. Digestion was incubated for 5 min at 37°C while shaking and stopped by adding 5 mM EDTA and then 8 ml pre-warmed RPMI-8226 was added. Cells were resuspended in PlasmaX™ + 2.5% human plasma + 1% Penicillin/Streptomycin + 150 IU/ml IL-2 and counted. 75,000 cells were seeded in a α-CD28 coated 96 well plate (final volume 150 µl). 194 nM melphalan was added or DMSO to vehicle control. Treatment was incubated for 3 days. For indirect treatment, MM.1s and RPMI-8226 were harvested and counted. 70,000 cells were seeded in a 24-well plate and cells were treated with either 194 nM melphalan (MM.1s) or 1.421 µM (RPMI-8226). Treatment was incubated for 3 days. Cells were harvested and spun down at 400 rcf for 5 min at 4°C. Supernatant was transferred on T cells 1:1 (v:v), MM cell line cells were washed once with DPBS and then lysed in 150 µl Extraction Buffer from the Arcturus® PicoRNA® isolation kit (Thermo Fisher Scientific, Cat. No. KIT0204). T cells were processed the same way like MM cell lines, except healthy donors were combined into one reaction to avoid interpatient heterogenity. RNA was isolated following the manufacturer's protocol. RNA quality and concentration were measured with NanoDrop™ One (Thermo Fisher Scientific, Cat. No. ND-ONE-W) and Invitrogen™ Qubit™ 4 fluorometer (Thermo Fisher Scientific, Cat. No. Q33238) with the Qubit™ RNA HS Assay Kit (Thermo Fisher Scientific, Cat. No. Q32855). 100 ng RNA was used as input in NEBNext® Single Cell/Low Input RNA Library Prep Kit for Illumina® (New England Biolabs, Cat. No. E6420L). Library was prepared following manufacturer's protocol, pooled and sequenced on an Illumina NextSeq2000 (paired-end, 2x 60 bp, 100 cycles).

##### *In vitro* mouse CD4$^+$ T cell isolation, melphalan stimulation and bulk RNAseq preparation

Spleens from wild-type C57BL/6 mice were harvested, and single cell suspensions were prepared. CD4+ T cells were isolated by positive selection using magnetic-activated cell sorting (MACS; L3T4 MicroBeads, Miltenyi Biotec, Cat. No. 130-117-043) with LS columns (Miltenyi Biotec, Cat. No. 130-042-401) and resuspended in Gibco™ Human plasma-like medium (HPLM; ThermoFisher, Cat. No. A4899101) supplemented with 10% FCS, 1% penicillin/streptomycin, and 1 µg/ml β-mercaptoethanol. Cells were counted and seeded in 24-well plates at a density of 1 x $10^6$ cells/ml (1 ml/well). Cells were then stimulated with 10 µM melphalan or DMSO for 24 or 72 hours. Finally, cells were collected, resuspended in 350 µl RLT buffer (provided in RNeasy Micro Kit, Qiagen, Cat. No. 74004) and stored at -80°C until further processing. RNA was then isolated with the RNeasy Micro Kit following the manufacturer's protocol. 19 ng of RNA was used as input into NEBNext® Single Cell/Low Input RNA Library Prep Kit for Illumina® and library was prepared following manufacturer's protocol, pooled and sequenced on an Illumina NextSeq2000 (paired-end, 2x 60 bp, 100 cycles).

##### Bulk RNAseq analysis

After performing quality control (QC) of the libraries using FASTQC, the raw reads were aligned to the human genome GRCh38 using the Rsubread 2.22.1 R package *(51)*. An expression matrix for the detected genes was then generated using the featureCounts function from the same package. Further analysis was performed using the DEseq2 1.48.2 R package *(52)*. Genes detected in less than three replicates and those with less than 10 reads were excluded for quality control purposes. Differentially expressed genes (DEGs) were identified between conditions by applying the DESeq function provided by DESeq2. Then, the DEGs were used to perform pathway enrichment analysis using clusterProfiler's v4.16.0 enrichment tools *(48)*. Lastly, pheatmap was used to create heatmaps, ggVennDiagram for generating venn diagrams and ggplot2 for the visualization of the data *(50, 53, 54)*.

**Metabolomics with liquid chromatography-mass spectrometry**

Plasma samples were thawed and sterile filtered (0.2 µm filter, Sarstedt, Cat. No. 83.1826.001) and frozen at -80°C. 5 µl of sample was diluted with 500 µl MeOH/H2O (80/20) containing external standard compounds (0.01 µM lamivudine and 1 µM each of D2-glucose, D4-succinate, D5-glycine and 15N-glutamate). After mixing and centrifugation (2 min at max. rcf in an Eppendorf centrifuge), the resulting supernatants were evaporated in a centrifugal evaporator. Samples were reconstituted in 150 µl of 5 mM NH4OAc in CH3CN/H2O (50/50, v/v). LC/MS analysis was performed on a Thermo Scientific Dionex Ultimate 3000 UHPLC system connected to a Q Exactive mass spectrometer (QE-MS) equipped with a HESI probe (ThermoFisher). Chromatographic separation was achieved by applying 3 µl sample on a XBridge Premier BEH Amide (100 × 2.1 mm, 2.5 µm) (Waters, Cat. No. 186009932) protected by a Supelco ColumnSaver particle filter (Merck, Cat. No. 55214-U and 55215-U) and a gradient of mobile phase A (5 mM NH4OAc in CH3CN/H2O (40/60, v/v)) and mobile phase B (5 mM NH4OAc in CH3CN/H2O (95/5, v/v)) maintaining a flow rate of 200 µl/min and a column temperature of 45°C. The LC gradient program was 100% mobile phase B for 2 min, followed by a linear decrease to 20% B within 23 min, maintaining 20% B for 21 min and returning to 100% B in 2 min, followed by 7 min 100% B for column equilibration before each injection. The eluent was directed to the QE-MS from 2.7 min to 46 min after sample application. Mass detection was conducted in alternating pos./neg. full scan mode (at 70k resolution, scan range m/z 69 - 1000, AGC target 1E6 and 200 ms max. injection time). HESI parameters: Sheath gas: 20, aux gas: 1, spray voltage: 3.0 kV, capillary temp.: 300°C, S-lens RF level: 50.0, aux gas heater temp.: 120°C. Manual curation and integration of chromatographic peaks were performed with TraceFinder 5.1 using a mass tolerance of +/- 2 mMUs.

**Analysis of Metabolomics**

Integrated peak areas were processed using a custom R-based analytical pipeline. Raw peak areas were normalized to the total metabolite content for each sample to ensure consistency across measurements. Metabolites with low-quality scores, as defined by the core facility, were excluded from further analysis. Features with missing values in more than two patient samples were removed, and the remaining missing values were imputed using a (minimum integrated peak area/5) approach. The resulting dataset was subsequently log2-transformed and scaled to generate a robust and standardized dataset for downstream analyses. Statistical analysis was performed with ordinary one-way ANOVA with multiple comparisons.

***In vitro* melphalan treatment**

PBMCs and BMMCs were thawed as described for flow cytometry. Cells were incubated overnight at 37°C in MM medium with a final concentration of 2 Mio cells/ml. Cells were counted again and 200,000 cells were seeded in 200 µl PlasmaX™ + 2.5% human plasma + 1% Penicillin/Streptomycin. Cells were treated with six different melphalan concentrations in a range from 194 nM to 9.7 µM (1xIC25 to 50xIC25 concentration). Samples were incubated for 3 days. Samples were then spun down at 532 rcf for 2 min at 4°C and then washed with DPBS. Staining was performed as described for flow cytometry with following antibodies: [α-human-CD3ε, AF700, clone: HIT3a, 1:300, Biolegend, Cat. No. 300324], [α-human-CD4, Brilliant Violet BV421, clone: Okt4, 1:300, Biolegend, Cat. No. 317434], [α-human-CD8a, APC, clone: RPA-T8, 1:300, ThermoFisher, Cat. No. 17-0088-42], [α-human-CD45, AF488, clone: 2D1, 1:50, Biolegend, Cat. No. 368535], [α-human-CD45RA, PerCP/Cy5.5, clone: HI100, 1:50, Biolegend, Cat. No. 304122], [α-human-CD62L, SB600, clone: DREG-56, 1:100, ThermoFisher, Cat. No. 63-0629-41]. In the end, samples were resuspended in FACS buffer and measured at the Attune™ NxT Flow Cytometer. Flow cytometry data was analyzed with FlowJo v10.8.1 (BD) and visualized with GraphPad Prism 9 (Insight Partners).

**Analysis of microarray data**

Microarray data of 57 samples from 29 elderly diagnosed plasma cell leukemia patients from the HOVON129 clinical trial was included in this study (GEO accession ID: GSE164703) *(55)*. Using the GEOquery 2.76.0 R package *(56)*, the expression profile was loaded and analyzed using the Bioconductor limma 3.64.3 R package *(57)*. Then differential expression analysis was performed on all genes that had a log2-transformed MAS5 expression value more than 5 in at least 10% of patients. The results were applied to perform pathway enrichment analysis using clusterProfiler's v4.16.0 enrichment tools *(48)*. Lastly, TF activity was inferred from normalized expression data using the decoupleR 2.14.0 R framework together with DoRothEA regulons loaded with dorothea R package with 1.20.0 version *(58, 59)*. SKY92 gene signature was collected and applied on the data for the risk stratification, by applying 0.827 threshold on the summed product of the multiplied values from normalized and standardized expression value of each of the signature genes with their assigned weighting factors *(16)*.

**CellROX staining**

500,000 cells of RPMI-8226 were seeded in 2 ml HPLM + 2.5% FBS + 1% Penicillin/Streptomycin and were treated with 10 µM melphalan or DMSO. Cells were incubated for 1h, 2h, 4h, 8h, 16h and 24h and harvested. Cells were stained with 5 µM CellROX™ deep red (ThermoFisher, Cat. No. C10422) for 30 min at 37°C. Afterwards, cells were washed 2 times with cold DPBS and then stained in 100 µl live/dead solution (Fixable viability dye, APC-eFluor780, 1:1000) for 15 min at 4°C. Cells were washed once with cold DPBS and then resuspended in FACS buffer and measured at the Attune™ NxT Flow Cytometer. Flow cytometry data was analyzed with FlowJo v10.8.1 (BD) and visualized with GraphPad Prism 9 (Insight Partners).

**Drug screening**

MM.1s and RPMI-8226 were collected and 4000 cells were seeded in 33 µl PlasmaX™ (CancerTools, Cat. No. 156371) + 2.5% FBS + 1% Penicillin/Streptomycin in a 384 well plate. Melphalan (Sigma-Aldrich, Cat. No. M2011) was added in 10 different concentrations in the range of 1 nM to 100 µM. Treatment was incubated for 3 days. Same volume of CellTiter-Glo® 3D Reagent (Promega, Cat. No. G9682) was added, 5 min slowly shaken (horizontally) and then 20 min further incubated at RT in the dark. Luminescence was measured for 500 ms with the TECAN Spark® Cyto plate reader (Tecan Group AG). Viability was calculated using following equating: Viability (%) = (Luminescence intensity treated/ luminescence DMSO control) * 100. Viability was further normalized to 1 nM concentration.

### Supplementary Figure 1

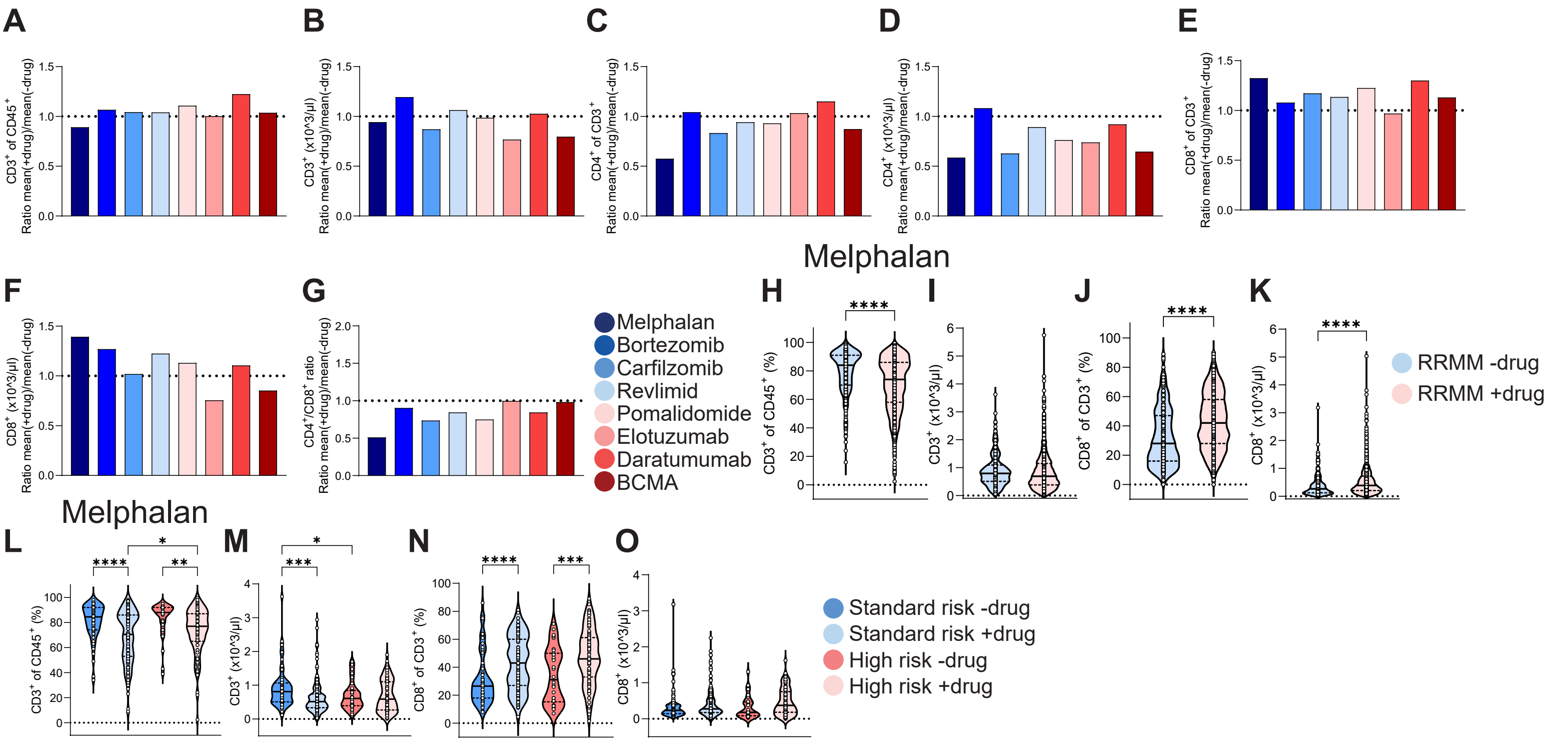

**Fig. S1. Melphalan has the highest impact on CD4+ T cells compared to other MM therapies.** (**A** to **G**) Barplot of effect of drugs on T cell populations in the PB of RRMM patients analyzed with flow cytometry. CD3+ of CD45+ (A), absolute number of CD3+ T cells (B), CD4+ of CD3+ (C), absolute number of CD4+ T cells (D), CD8+ of CD3+ (E), absolute number of CD8+ T cells (F), and CD4+/CD8+ T cell ratio (G). (**H** to **K**) Quantification of flow cytometry from PBMC of RRMM patients in dependency of melphalan treatment. Percentage of CD3+ of CD45+ (-drug: n=322; +drug: n=1357) (H), absolute number of CD3+ T cells (-drug n=281; +drug n=1169) (I), percentage of CD8+ of CD3+ (-drug: n=318; +drug: n=1301) (J), and absolute number of CD8+ T cells (-drug: n=279; +drug: n=1120) (K). (**L** to **O**) Quantification of flow cytometry from PB of RRMM patients in dependency of melphalan treatment and risk status. Percentage of CD3+ of CD45+ (SR -drug n=64; SR +drug n=156; HR -drug n=34; HR +drug n=100) (L), absolute number of CD3+ T cells (SR -drug n=61; SR +drug n=135; HR -drug n=33; HR +drug n=80) (M), percentage of CD8+ of CD3+ (SR -drug n=64; SR +drug n=153; HR -drug n=34; HR +drug n=98) (N), and absolute number of CD8+ T cells (SR -drug n=61; SR +drug n=134; HR -drug n=33; HR +drug n=78) (O). * $p < 0.05$, ** $p < 0.01$, *** $p < 0.001$, **** $p < 0.0001$.

### Supplementary Figure 2

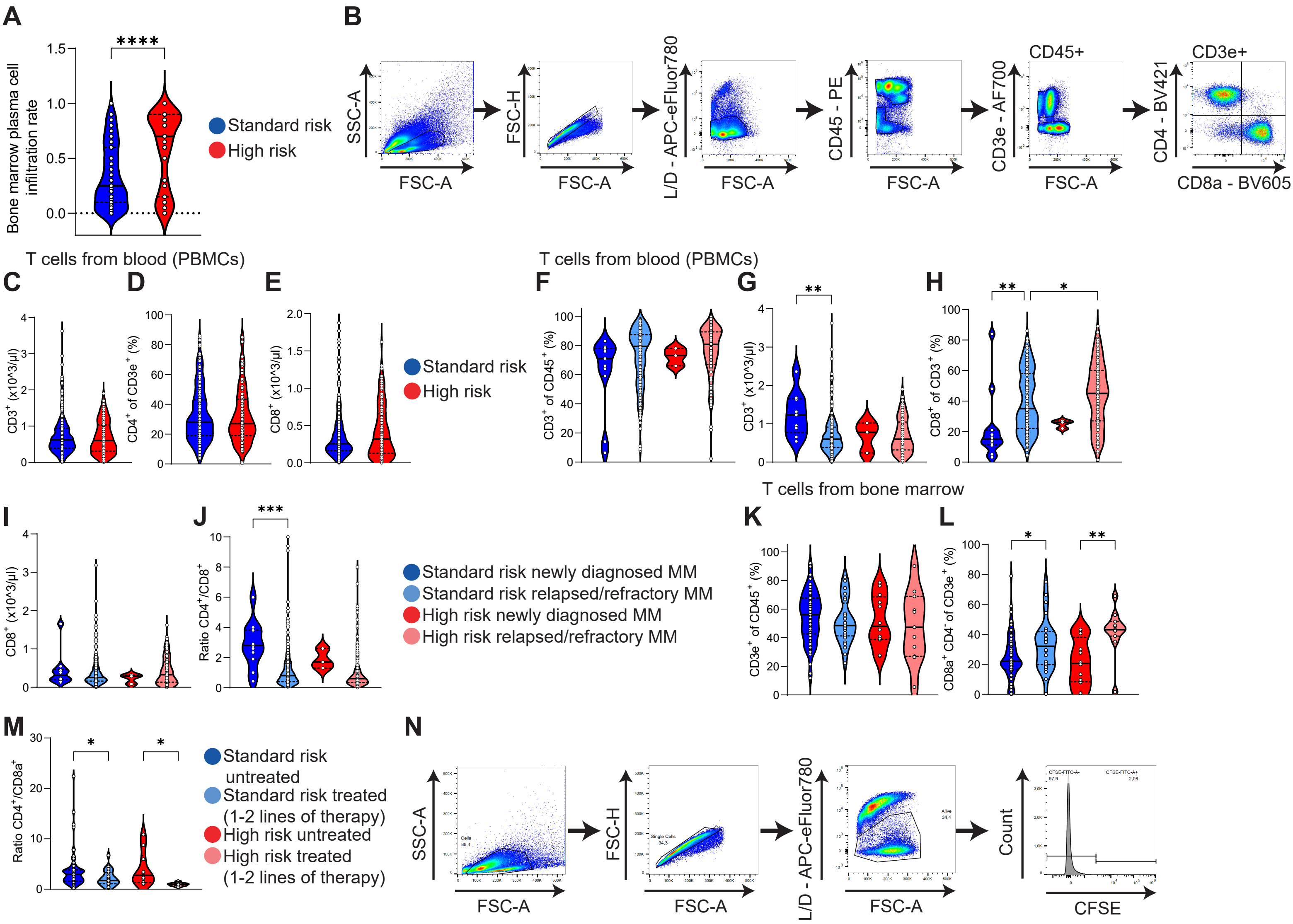

**Fig. S2. Treatment induces an increase in the proportion of CD8+ T cells in both risk groups.** (**A**) BM PC infiltration rate of high-risk and standard-risk patients (SR n=115; HR n=45). (**B**) Gating strategy for flow cytometry analysis. (**C** to **E**) Quantification of flow cytometry from PB of RRMM patients in dependency of risk status. Absolute number of CD3+ (SR n=206; HR n=116) (C), proportion of CD4+ of CD3+ T cells (SR n=232; HR n=136) (D), and absolute number of CD8+ T cells (SR n=206; HR n=114) (E). (**F** to **J**) Quantification of flow cytometry from PB of RRMM patients in dependency of risk status and treatment status. Proportion of CD3+ T cells of CD45+ (SR NDMM n=11; SR RRMM n=221; HR NDMM n=3; HR RRMM n=134) (F), absolute number of CD3+ (SR NDMM n=9; SR RRMM n=197; HR NDMM n=3; HR RRMM n=113) (G), proportion of CD8+ of CD3+ T cells (SR NDMM n=12; SR RRMM n=218; HR NDMM n=3; HR RRMM n=132) (H), absolute number of CD8+ T cells (SR NDMM n=10; SR RRMM n=196; HR NDMM n=3; HR RRMM n=111) (I), and CD4+/CD8+ T cell ratio (SR NDMM n=12; SR RRMM n=217; HR NDMM n=3; HR RRMM n=129) (J). (**K** to **M**) Quantification of flow cytometry from BM of RRMM patients in dependency of risk status and treatment status. Proportion of CD3e+ T cells of CD45+ (SR untreated n=72; SR 1-2 lines of therapy n=28; HR untreated n=12; HR 1-2 lines of therapy n=11) (K), proportion of CD8a+ of CD3+ T cells (SR untreated n=73; SR 1-2 lines of therapy n=28; HR untreated n=12; HR 1-2 lines of therapy n=11) (L), and CD4+/CD8+ T cell ratio (SR untreated n=73; SR 1-2 lines of therapy n=28; HR untreated n=12; HR 1-2 lines of therapy n=11) (M). (**N**) Gating strategy of in vitro cytotoxicity assay.

### Supplementary Figure 3

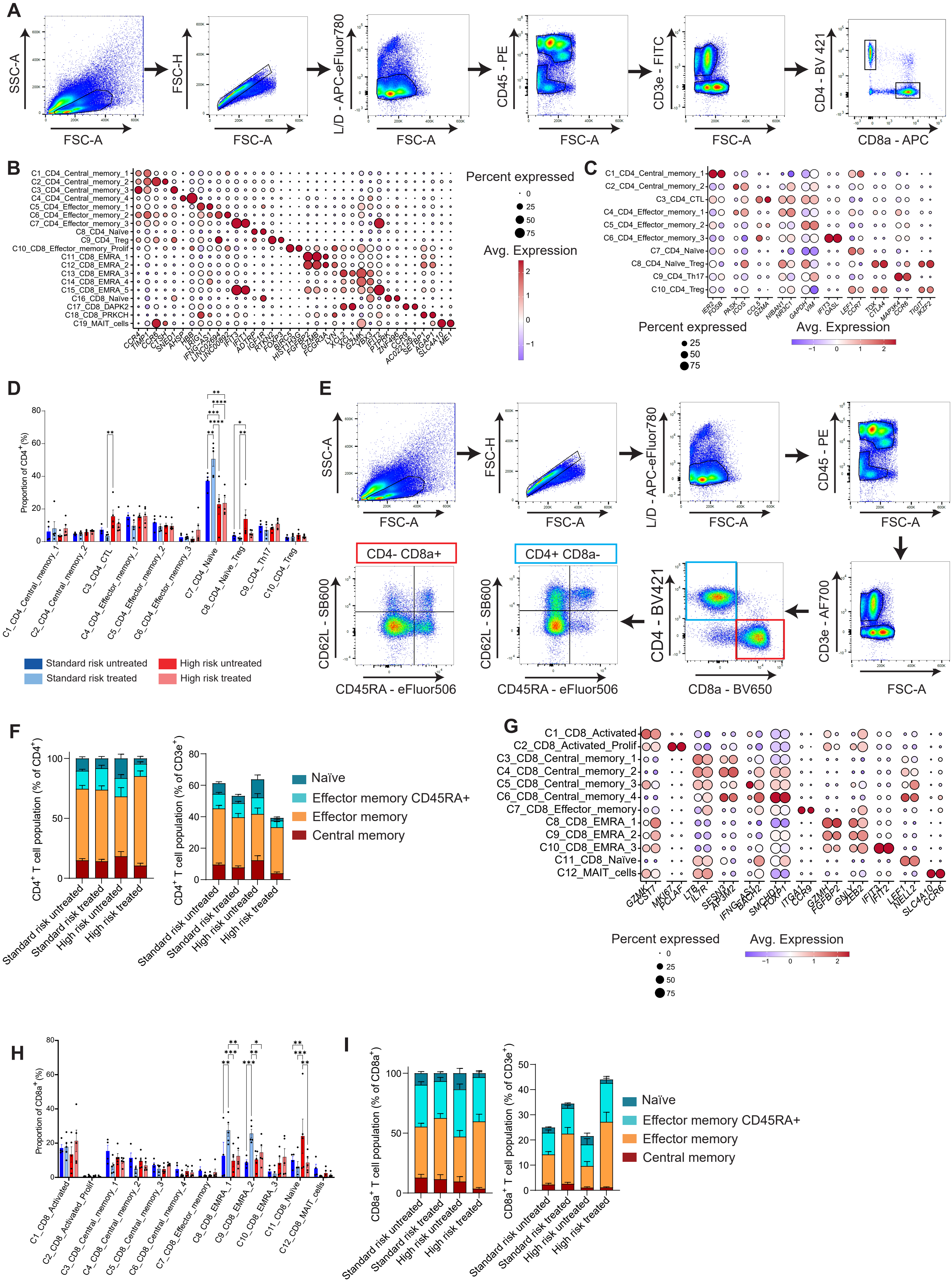

**Fig. S3. CD8+ effector memory CD45RA+ T cells are enriched after treatment.** (**A**) Gating strategy for sorting for scRNAseq. (**B**) Average expression of top 2 marker genes per cluster from scRNAseq data in dotplot. (**C**) Dotplot showing average expression of top 2 marker genes per cluster from the separated CD4+ T cell compartment. (**D**) Barplot showing proportion of cells per patient in each of the CD4+ T cell population. Ordinary two-way ANOVA with Dunnett's test as a Post-hoc test was used for statistical analysis. (**E**) Gating strategy for T cell subset quantification. (**F**) Quantification of flow cytometry of T cell subsets from the BM. Proportion of CD4+ subsets from CD4+ T cells (left panel) and proportion of CD4+ T cell subsets of CD3e+ T cells (right panel). Sea green represents naïve T cells, light blue represents effector memory CD45RA+ T cells, orange represents effector memory T cells, and red represents central memory T cells. (**G**) Dotplot showing average expression of top 2 marker genes per cluster from the separated CD8+ T cell compartment. (**H**) Barplot showing proportion of cells per patient in each of the CD8+ T cell population. Ordinary two-way ANOVA with Dunnett's test as a Post-hoc test was used for statistical analysis. (**I**) Quantification of flow cytometry of T cell subsets from the BM. Proportion of CD8+ subsets from CD8+ T cells (left panel) and proportion of CD8+ T cell subsets of CD3e+ T cells (right panel). Sea green represents naïve T cells, light blue represents effector memory CD45RA+ T cells, orange represents effector memory T cells, and red represents central memory T cells. * $p < 0.05$, ** $p < 0.01$, *** $p < 0.001$, **** $p < 0.0001$.

Supplementary Figure 4

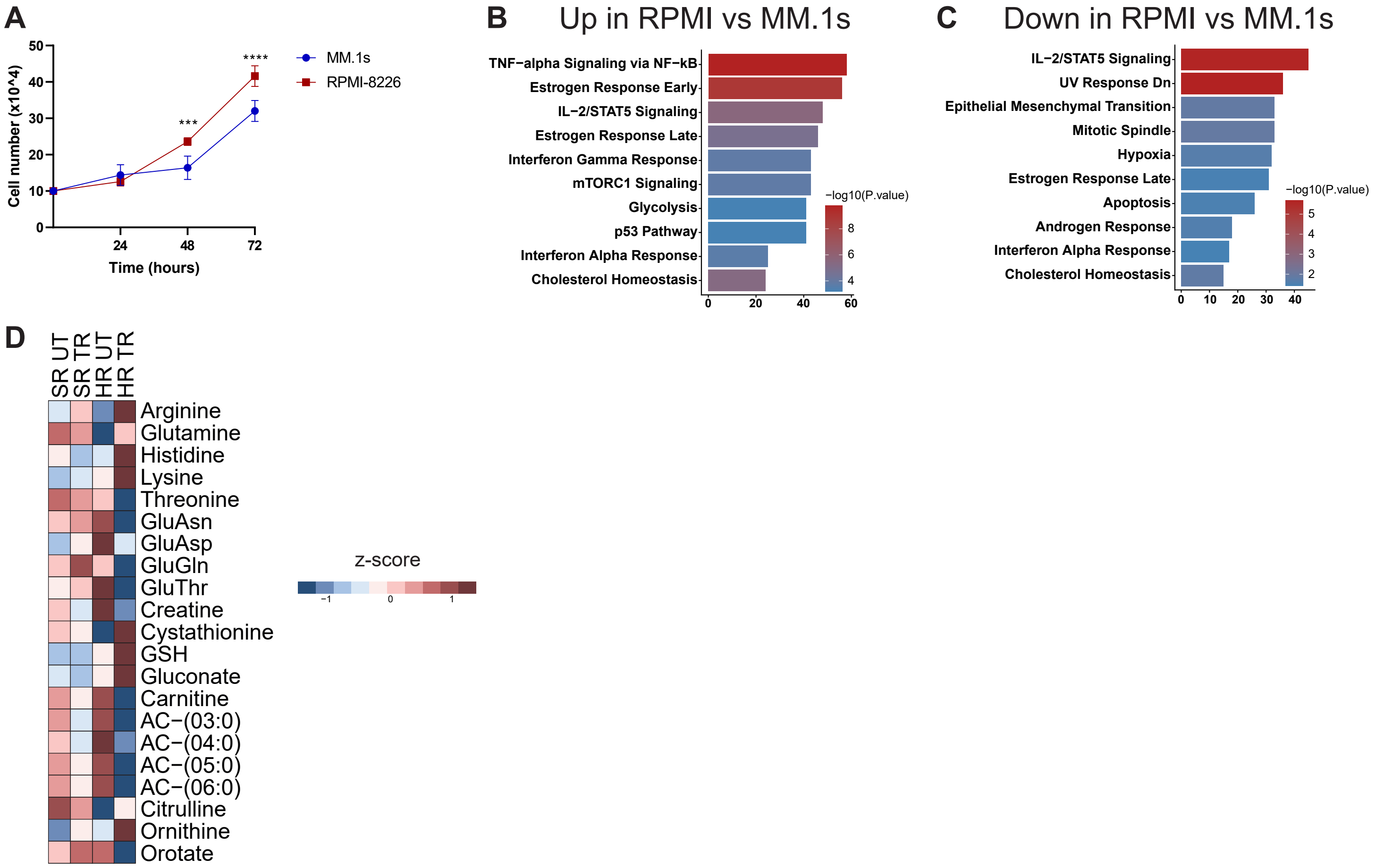

**Fig. S4. RPMI-8226 represents a more high-risk cell line.** (**A**) Cell doubling experiment of MM cell lines MM.1s and RPMI-8226. Blue indicates MM.1s and red indicates RPMI-8226. (**B**) Hallmark gene sets analysis of upregulated genes in RPMI-8226 compared to MM.1s. (**C**) Hallmark gene sets analysis of downregulated genes in RPMI-8226 compared to MM.1s. (**D**) Heatmap of z-scores from water-soluble metabolites from the BM plasma of SR and HR patients of all significant metabolites.

### Supplementary Figure 5

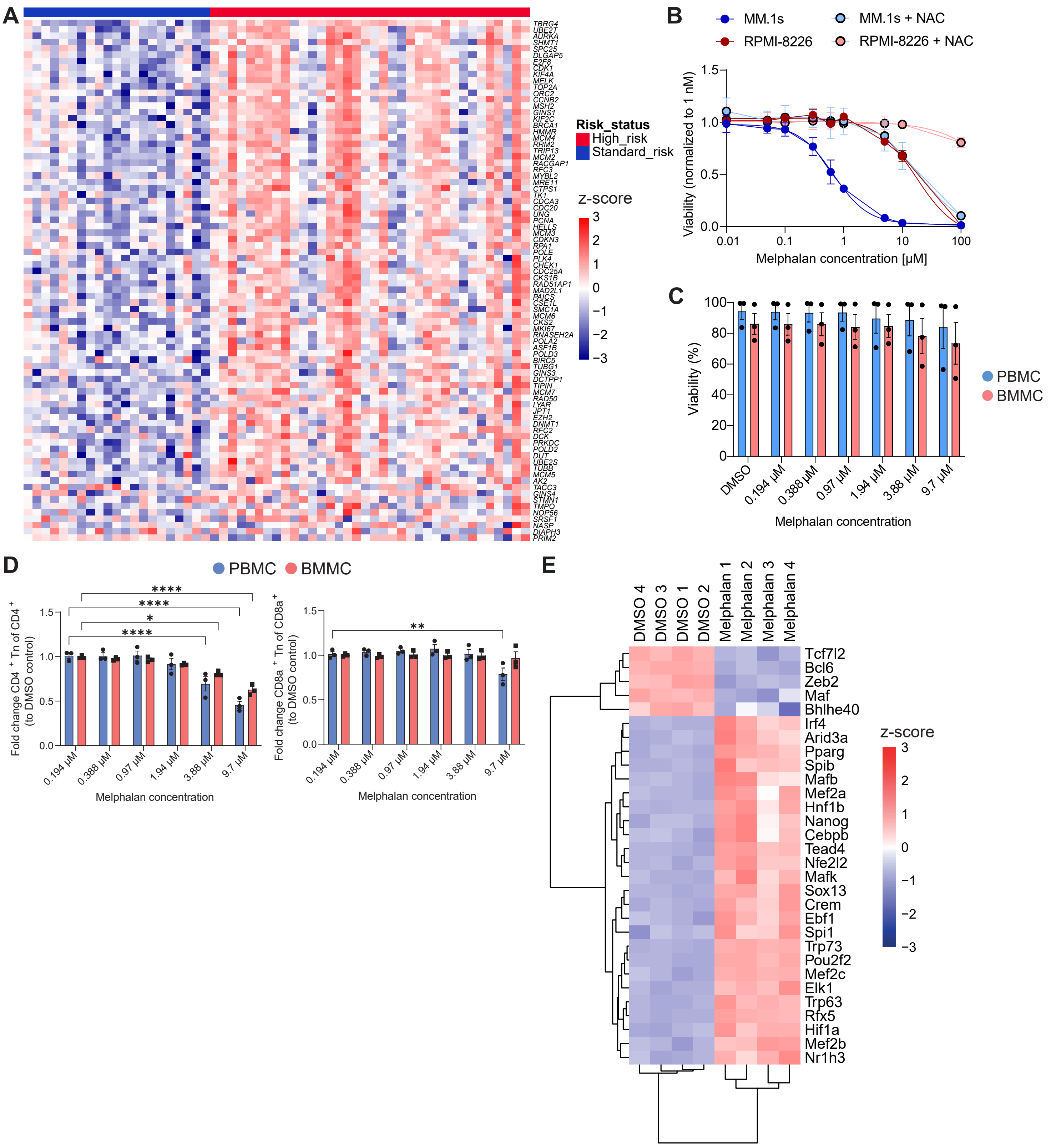

**Fig. S5. In vitro melphalan treatment depletes CD4+ naïve T cells.** (**A**) Heatmap of genes linked with E2F genes pathway from Fig. 5G. (**B**) Viability of cells treated with different melphalan concentration and +/- 1 mM NAC. MM.1s in blue and RPMI-8226 in red. NAC treated cells are indicated by black border and lighter colors. Graphs with only melphalan treatment were already shown in Fig. 4A. (**C**) *In vitro* treatment of PB and BM mononuclear cells with melphalan (n=3 for each group). Plotted is the viability of all cells. (**D**) *In vitro* treatment of PB and BM mononuclear cells with melphalan (n=3 for each group). Left panel: CD4+ naïve T cells, right panel: CD8a+ naïve T cells. Ordinary two-way ANOVA with Dunnett's test as a Post-hoc test was used for statistical analysis. (**E**) Heatmap showing variable TF activities across the samples. * $p < 0.05$, ** $p < 0.01$, *** $p < 0.001$, **** $p < 0.0001$.

Supp Table 1

| **PBMC RRMM** | |
|---|---|
| Age (mean [range]) | 65 [32-86] |
| Gender | |
| female | 110 |
| male | 242 |
| Lines of therapy | |
| 1 | 701 |
| 2 | 287 |
| 3 | 203 |
| 4 | 176 |
| ≥5 | 320 |
| Risk status | |
| Standard risk | 33 |
| High risk | 26 |
| N.a. | 295 |

Supp Table 2

| **PBMC** | (HR vs SR) | |
|---|---|---|
| | SR | HR |
| Age (mean [range]) | 66 [50-86] | 63 [31-83] |
| Gender | | |
| female | 12 (28%) | 11 (39%) |
| male | 31 (72%) | 17 (61%) |
| Lines of therapy | | |
| 0 | 13 | 3 |
| 1 | 147 | 39 |
| 2 | 21 | 11 |
| 3 | 13 | 25 |
| 4 | 6 | 14 |
| ≥5 | 34 | 45 |

Supp Table 3

| **BMMC** | | |
|---|---|---|
| | SR | HR |
| Age (mean [range]) | 64 [32-86] | 61 [33-82] |
| Gender | | |
| female | 52 (44%) | 20 (44%) |
| male | 65 (56%) | 25 (56%) |
| Lines of therapy | | |
| 0 | 72 | 12 |
| 1 | 18 | 8 |
| 2 | 10 | 3 |
| 3 | 4 | 4 |
| 4 | 5 | 3 |
| ≥5 | 8 | 15 |
| BMPCIR (mean [range]) | 0.34 [0-1] | 0.54 [0-1] |

**Supplementary Table 1** Information of all relapsed of refractory multiple myeloma patients from peripheral blood mononuclear cells samples.

**Supplementary Table 2** Information of relapsed of refractory multiple myeloma patients from peripheral blood mononuclear cells samples with either standard-risk (SR) or high-risk (HR) disease.

**Supplementary Table 3** Information of multiple myeloma patients from bone marrow mononuclear cells samples with either standard-risk (SR) or high-risk (HR) disease.
